# Two enhancer binding proteins activate σ^54^-dependent transcription of a quorum regulatory RNA in a bacterial symbiont

**DOI:** 10.1101/2022.04.27.489697

**Authors:** Ericka D. Surrett, Kirsten R. Guckes, Shyan Cousins, Terry B. Ruskoski, Andrew G. Cecere, Denise A. Ludvik, C. Denise Okafor, Mark J. Mandel, Tim I. Miyashiro

## Abstract

To colonize a host, bacteria depend on an ensemble of signaling systems to convert information about the various environments encountered within the host into specific cellular activities. How these signaling systems coordinate transitions between cellular states *in vivo* remains poorly understood. To address this knowledge gap, we investigated how the bacterial symbiont *Vibrio fischeri* initially colonizes the light organ of the Hawaiian bobtail squid *Euprymna scolopes*. Previous work has shown that the small RNA Qrr1, which is a regulatory component of the quorum-sensing system in *V. fischeri*, promotes host colonization. Here, we report that transcriptional activation of Qrr1 is repressed by the regulator BinK, which inhibits cellular aggregation by *V. fischeri* prior to light organ entry. We show that Qrr1 expression depends on the alternative sigma factor σ^54^ and the transcription factors LuxO and SypG, which function together as an OR logic gate, thereby ensuring Qrr1 is expressed during colonization. Finally, we provide evidence that this regulatory mechanism is widespread throughout the *Vibrionaceae* family. Together, our work reveals how coordination between the signaling pathways underlying aggregation and quorum sensing promotes host colonization, which provides insight into how integration among signaling systems facilitates complex processes in bacteria.

## Introduction

The overall fitness of an animal often depends on the activities of bacteria that are localized to certain anatomical sites of the host. In many cases, these bacteria are horizontally transmitted among hosts, which means that they are first shed into a reservoir prior to colonizing a new host. The environmental conditions associated with the reservoir are typically different than those encountered on or within a host. Therefore, to properly acclimate to an environment, bacteria depend on signal transduction systems that coordinate cellular physiology in response to a vast array of environmental signals and cues. How these signaling pathways facilitate the cellular activities that are pertinent to the complex environments encountered during host colonization remains unclear for most bacteria. Focusing on the connections between different signaling pathways has the potential to fill this knowledge gap and provide insight into how bacteria transition from one environment to another.

The bioluminescent bacterium *Vibrio fischeri* (also known as *Aliivibrio fischeri*) is a notable example of a bacterium that depends on multiple signaling systems to establish and maintain association with a host (1–3). While a variety of marine animals serve as hosts for *V. fischeri*, the Hawaiian bobtail squid *Euprymna scolopes* is by far the best characterized, and this host-microbe association has emerged as a powerful system to model how signaling systems function in a natural host environment. From a specialized light organ located within the ventral side of the mantle, populations of *V. fischeri* emit bioluminescence that camouflage the host when viewed from below (4). Because *V. fischeri* grows on host-derived compounds within the light organ (5), the association is considered a mutualistic symbiosis, in which each taxon benefits from their long-term and intimate interactions. The symbiosis is initially established after juvenile squid are exposed to seawater containing *V. fischeri* cells (6), which enables bacterial mutants to be assessed for their ability to establish symbiosis, *i.e.*, to colonize, grow, and produce bioluminescence within the light organ.

The light-producing luciferase enzyme is encoded within the *lux* operon, which is transcribed when signaling by the LuxI/LuxR quorum-sensing system occurs (1) (Fig. 1). Quorum sensing describes the phenomenon when bacteria synthesize, detect, and respond to small signaling molecules called autoinducers (7, 8). Mutants for either LuxI (autoinducer synthase) or LuxR (autoinducer receptor) fail to produce bioluminescence *in vivo* (9), which illustrates the significance of quorum sensing for the symbiosis to be established. In addition to the LuxI/LuxR system, two other quorum-sensing systems (AinS/AinR and LuxS/LuxPQ) affect bioluminescence production by indirectly regulating transcription of the *lux* operon (1) (Fig. 1). Under conditions of low autoinducer concentrations, either AinR or LuxPQ can trigger a phosphorelay that results in phosphorylation of the transcription factor LuxO (10, 11). In conjunction with the alternative sigma factor σ^54^, LuxO activates the transcription of the small quorum regulatory RNA Qrr1 (10, 12), which lowers the ability of *V. fischeri* to enhance bioluminescence production (Fig. 1). In contrast to the critical role that the LuxI/LuxR system has on establishing symbiosis, the impact of signaling by these other quorum-sensing systems is more nuanced, with knockout mutants for specific pathway components exhibiting symbiosis-related phenotypes that are observable only when introduced to juvenile squid as an inoculum mixed with another strain type (reviewed in (2)). For instance, a Δ*qrr1* mutant can establish a light-organ symbiosis with bacterial abundance and bioluminescence emission levels that are indistinguishable from squid colonized with the wild-type strain (10). However, when juvenile squid are exposed to an inoculum evenly mixed with Δ*qrr1* mutant and wild-type strains, they later feature light organs containing fewer Δ*qrr1* cells than wild-type cells (10), which suggests that the expression of Qrr1 provides an advantage for *V. fischeri* to establish symbiosis when other potential founder cells are also present.

**Figure 1:**
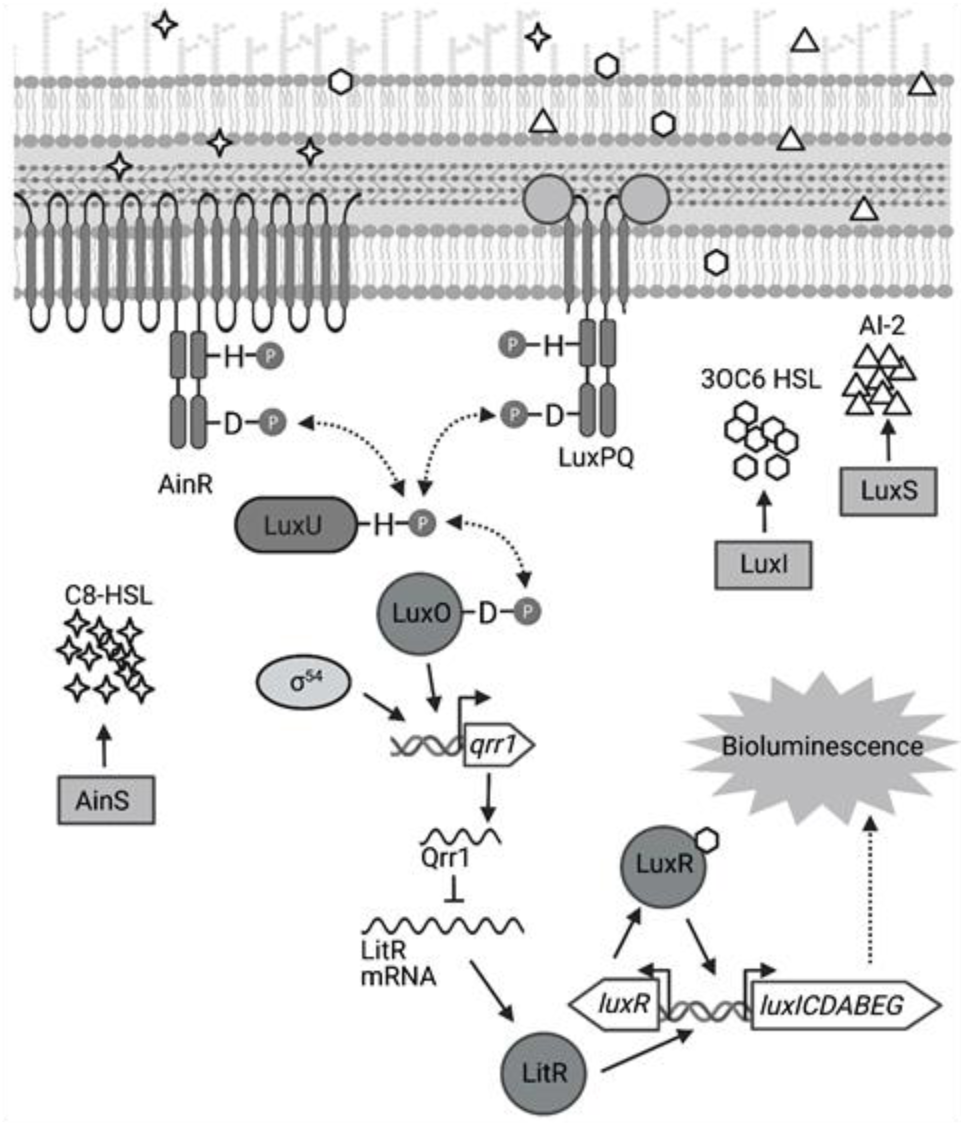
Signal transduction network underlying quorum sensing in *Vibrio fischeri*. Quorum sensing in *V. fischeri* depends on autoinducers 3-oxo-C6 HSL (3OC6 HSL), AI-2, and C8-HSL, which are synthesized by LuxI, LuxS, and AinS, respectively. Interaction of C8-HSL or AI- 2 with their cognate sensors (AinR and LuxPQ, respectively), results in lower levels of phosphorylated LuxO. Phosphorylated LuxO promotes σ^54^-dependent transcription of *qrr1*, which encodes the sRNA Qrr1. Qrr1 post-transcriptionally represses LitR, which is a positive regulator of *luxR*. Consequently, quorum sensing inhibits Qrr1 expression, thereby promoting bioluminescence production.

The primary structure of LuxO features an N-terminal regulatory domain, a central catalytic domain, and a C-terminal DNA-binding domain that define this transcription factor as a Group I bacterial enhancer binding protein (bEBP) (13). As extensively reviewed elsewhere (13, 14), bEBPs bind upstream of σ^54^-dependent promoters and hydrolyze ATP to induce the conformational changes within the RNA polymerase/σ^54^/promoter complex that facilitate transcription initiation. Mechanistic studies in other *Vibrionaceae* have shown that the ATPase activity of LuxO is controlled by its N-terminal regulatory domain (15), which consists of a REC domain that participates in the phosphorelay (16) (Fig. 1). In its unphosphorylated form, LuxO is inactive, with a 20-residue linker that connects the regulatory and catalytic domains occupying the active site within the catalytic domain to block nucleotide binding (15). The linker is a structural feature reportedly unique to LuxO, and its position within the active site is stabilized by hydrogen bonds with the regulatory and catalytic domains (15). This linker model is also supported for the LuxO homolog of *V. fischeri*—V114 is a residue within the regulatory domain that is predicted to interact with the linker region, and its substitution with either alanine or glycine results in a variant of LuxO with elevated activity (12). Phosphorylation of an aspartate conserved among REC domains (D55 in the LuxO homolog of *V. fischeri*) is predicted to displace the linker (17), which enables activation of LuxO and transcriptional initiation of the *qrr1* promoter (*P_qrr1_*). Phosphorylation of LuxO occurs when the histidine kinases that serve as quorum-sensing receptors are unbound with ligand (11) (Fig. 1); consequently, conditions of low cell density result in LuxO activity and transcriptional activation of *P_qrr1_* (10). As the population grows, higher levels of the respective autoinducer ligands promote LuxO de-phosphorylation, thereby lowering Qrr1 expression and permitting enhanced bioluminescence production (Fig. 1).

Despite these advances in understanding the molecular mechanisms by which quorum-sensing systems regulate transcriptional activity of *P_qrr1_* in *V. fischeri* (10, 18), how Qrr1 expression is controlled during symbiosis establishment remains unclear. For instance, prior to entering the light organ, bacterial cells are collected from the environment and form aggregates that are densely packed (3, 19). *A priori*, such cellular arrangements are predicted to engage in quorum sensing and lower transcriptional activation of *P_qrr1_* (Fig. 1), which would seemingly prevent cells from expressing Qrr1 to gain an advantage in host colonization. Here, we report a regulatory mechanism that enables *V. fischeri* to avoid this predicament. In particular, we reveal that the signaling pathways associated with aggregation and quorum sensing are connected in *V. fischeri*, and we demonstrate that this connection contributes to host colonization.

Genetic analysis shows that σ^54^-dependent transcription of *P_qrr1_* can be activated by two distinct bEBPs that depend on overlapping *cis* regulatory elements, thereby resulting in a gene regulation module that resembles an OR logic gate, in which activation of either bEBP results in Qrr1 expression. Bioinformatic analysis suggests the potential for dual bEBP activation of Qrrs in approximately half of the other clades of the *Vibrionaceae* family, which suggests that this regulatory mechanism is widespread among biomedically and ecologically important taxa.

## Results

### BinK inhibits transcriptional activation of Qrr1

In *V. fischeri*, transcriptional activity of *P_qrr1_* lowers in response to the autoinducer C8 HSL (11), which indicates that quorum sensing attenuates Qrr1 expression. Consequently, the high cell density associated with colonies leads to low *P_qrr1_* transcriptional activity. Previously, we described a screen designed to identify genetic factors that inhibit *P_qrr1_* activity within colonies (20). More specifically, the screen had been performed by introducing a GFP-based, transcriptional reporter for *P_qrr1_* (*P_qrr1_*::*gfp*) into a Tn*5*-mutant library derived from wild-type strain ES114, selecting for conjugants by plating cells onto solid rich medium, and screening the resulting colonies for increased GFP fluorescence. One mutant resulting from the screen contains a transposon insertion within the gene *binK* (*VF_A0360*), which encodes the hybrid histidine kinase BinK (21) (Fig. 2A). To validate that the disruption of *binK* conferred increased *P_qrr1_* activity, we assessed the *P_qrr1_*::*gfp* reporter in a Δ*binK* mutant that was previously reported (21). When grown on solid medium to high cell density, the Δ*binK* mutant exhibited 3.7-fold higher levels of GFP fluorescence relative to WT (Fig. 2B), which suggests that *P_qrr1_* is transcriptionally active in cells lacking BinK. Wild-type levels of GFP fluorescence were observed in a Δ*binK* mutant expressing *binK in trans* (Fig. 2B), demonstrating genetic complementation. Together, these data suggest that conditions of high cell density fail to lower Qrr1 expression in cells lacking BinK.

**Figure 2:**
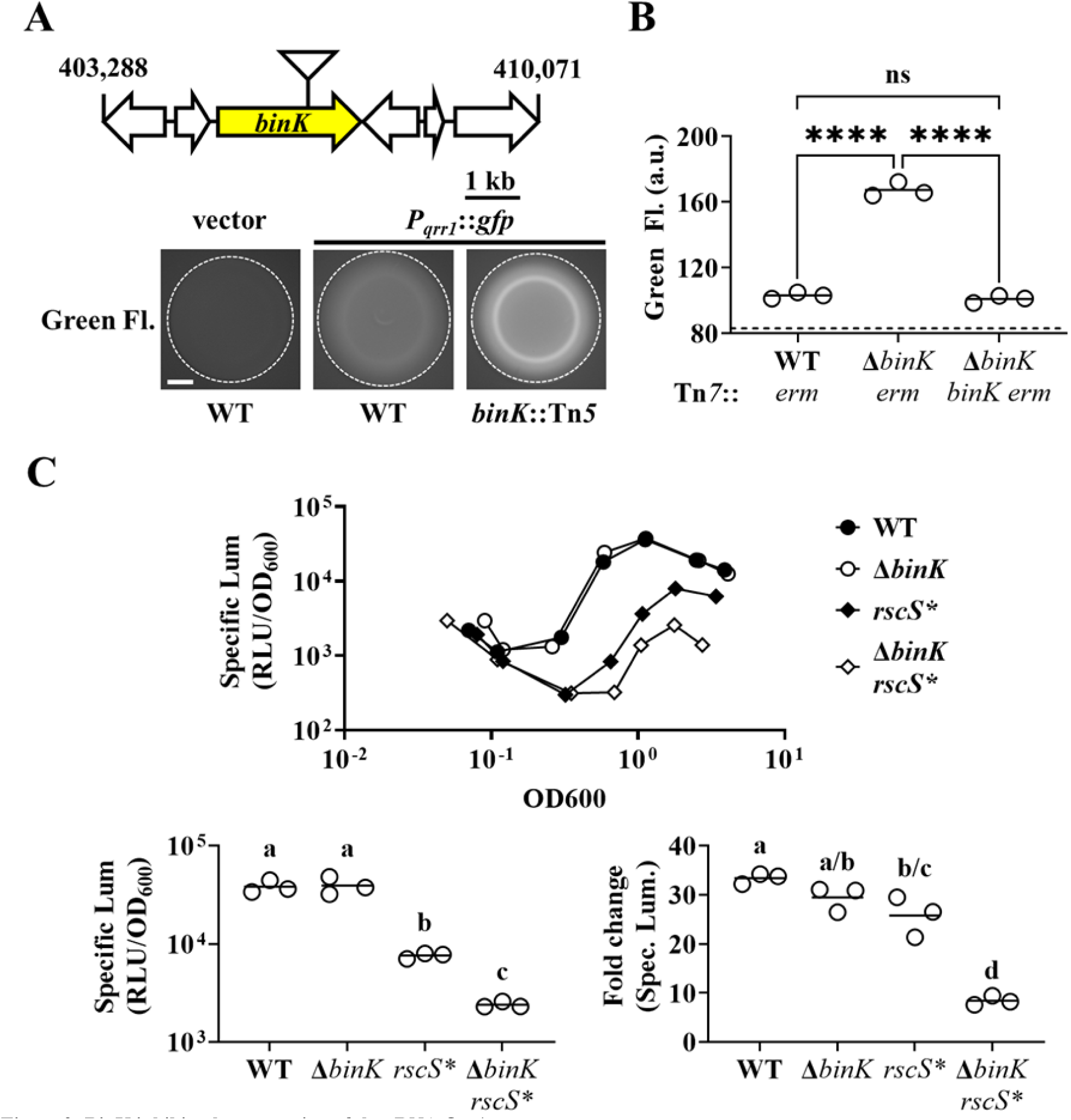
BinK inhibits the expression of the sRNA Qrr1. **(A)** *Top*, Tn*5* insertion location within *VF_A0360* (*binK*). *Below*, Green fluorescence images associated with ES114 (WT) and DRO22 (*binK*::Tn*5*) harboring pTM268 (*P_qrr1_*::*gfp*) or pVSV105 (vector). Dotted circle indicates border of the spot of bacterial growth resulting from placing a cell suspension on the surface of solid rich medium and incubating the sample at 28°C for 24 hours. Scale bar = 1 mm. **(B)** Green fluorescence levels of TIM313 (WT Tn*7*::*erm*), MJM2481 (Δ*binK* Tn*7*::*erm*), and TIM412 (Δ*binK* Tn*7*::[*binK erm*]) harboring pTM268. Point = green fluorescence of a spot (N = 3), bar = group mean. Dotted line = autofluorescence cutoff. One-way ANOVA (F_2,6_ = 466.9, *p* < 0.0001); Tukey’s *post-hoc* test with *p*-values corrected for multiple comparisons (n.s. = not significant, **** = *p* < 0.0001). **(C)** *Top*, Bioluminescence assay of ES114 (WT), MJM2251 (Δ*binK*), MJM1198 (*rscS\**), and MJM2255 (Δ*binK rscS\**). Point = specific luminescence (RLU/OD_600_) of indicated strain at the indicated turbidity (OD_600_). Shown are points derived from a representative culture (N = 3). *Bottom left*, Point = peak specific bioluminescence derived from each culture, with bar = group mean. One-way ANOVA on log- transformed data (F_3,8_ = 308.6, p < 0.0001); Tukey’s *post-hoc* test (same letters = not significant; different letters = *p* < 0.0001). *Bottom right*, Point = fold change in specific bioluminescence from min to max. One-way ANOVA (F3,8 = 308.6, p < 0.0001); Tukey’s *post-hoc* test (same letters = not significant; a-b = p < 0.05, c-d = p < 0.001, a-c & c-d = *p* < 0.0001). Experimental trials: 2.

In *V. fischeri*, Qrr1 post-transcriptionally represses the expression of LitR, which is a transcription factor that enhances transcription of the *lux* operon, so cells expressing Qrr1 produce low levels of bioluminescence (10). However, conditions of high cell density attenuate Qrr1 expression, which enables increased LitR expression and consequently higher cellular bioluminescence as cell abundance increases. Therefore, the observation of increased expression of Qrr1 associated with the Δ*binK* mutant prompted us to investigate bioluminescence production throughout growth in culture. The Δ*binK* mutant produces wild-type levels of bioluminescence, including when the bioluminescence emission per cell unit (specific luminescence) amplifies during exponential growth (Fig. 2C), which seemingly suggests that BinK has no impact on how quorum sensing regulates bioluminescence production. However, when originally assessed in a biofilm assay, the Δ*binK* mutant phenocopied the wild-type strain unless the histidine kinase RscS was overexpressed, which revealed that BinK inhibits biofilm formation (21). Therefore, we hypothesized that phenotypes associated with the Δ*binK* allele are similarly masked in bioluminescence assays. To test this hypothesis, we measured bioluminescence production of strains harboring the *rscS\** allele, which overexpresses RscS (22). Relative to the wild-type strain, the *rscS*\* mutant exhibited a specific bioluminescence profile with a lower peak and less amplification (Fig. 2C). The specific bioluminescence profile of the Δ*binK rscS\** mutant featured an even lower peak and lower amplification (Fig. 2C), which suggests that RscS overexpression reveals the ability of BinK to inhibit bioluminescence production. Taken together, these results provide evidence that the cellular physiology of Δ*binK* mutant can inhibit bioluminescence production and amplification under conditions of high cell density, which is consistent with elevated Qrr1 levels.

### Enhanced crypt colonization by the Δ*binK* mutant is independent of Qrr1

Qrr1 and BinK are significant factors in the life cycle of *V. fischeri* because they each impact how *fischeri* cells initially establish symbiosis with *E. scolopes*. BinK inhibits the aggregation that occurs among environmental *V. fischeri* cells collected by the light organ, such that cells lacking BinK form large aggregates prior to light organ entry (21, 23). In addition, animals exposed to an inoculum mixed evenly with a Δ*binK* mutant and its wild-type parental strain result in more Δ*binK* cells than wild-type cells within their light organs (21), which suggests that BinK inhibits the ability of a cell to establish symbiosis in the context of other colonizing bacteria. In contrast, Qrr1 provides an advantage to *V. fischeri* when establishing symbiosis in the presence of other cells, as squid exposed to an inoculum mixed evenly with a Δ*qrr1* mutant and its wild-type parental strain lead to fewer Δ*qrr1* cells than wild-type cells within colonized animals (10). Consequently, the discovery that *P_qrr1_* expression is elevated within a Δ*binK* mutant led us to investigate whether this regulatory connection impacts how *V. fischeri* establishes symbiosis, particularly in the context of competition.

Upon symbiosis establishment, the light organ contains up to six independent populations of *V. fischeri*, with each population housed within an epithelium-lined crypt space (24). Because the isolation of colony-forming units (CFUs) requires tissue homogenization, approaches based on counting CFUs to quantify cellular abundance *in vivo* inherently disrupt the location of the strains within the light organ, thereby precluding insight that can be deduced from this knowledge. Therefore, we first determined where the Δ*qrr1* mutant and a wild-type competitor strain reside within the light organ by differentially labeling each strain type with fluorescent proteins and assessing their location within host tissue by fluorescence microscopy (25) (Fig. 3A). As expected, most light organs contained populations in several crypt spaces (Figs. 3B-C), which indicated that multiple colonization events had occurred within each animal. The majority of crypt spaces each contained only one strain type (Fig. 3B), which is consistent with most populations arising from only 1-2 cells that enter the corresponding crypt spaces (26). Few crypt spaces harbored the Δ*qrr1* mutant (Fig. 3B-C), which suggests that the majority of populations were founded by wild-type cells. In contrast, when the inoculum contained an equal mix of differentially labeled wild-type cells, no difference was observed in the number of crypt spaces colonized by YFP- or CFP-labeled strains (Figs. 3B-C). Consequently, these results suggest that the competitive defect of the Δ*qrr1* mutant reported previously is due to fewer crypt spaces being initially accessed by the mutant.

**Figure 3:**
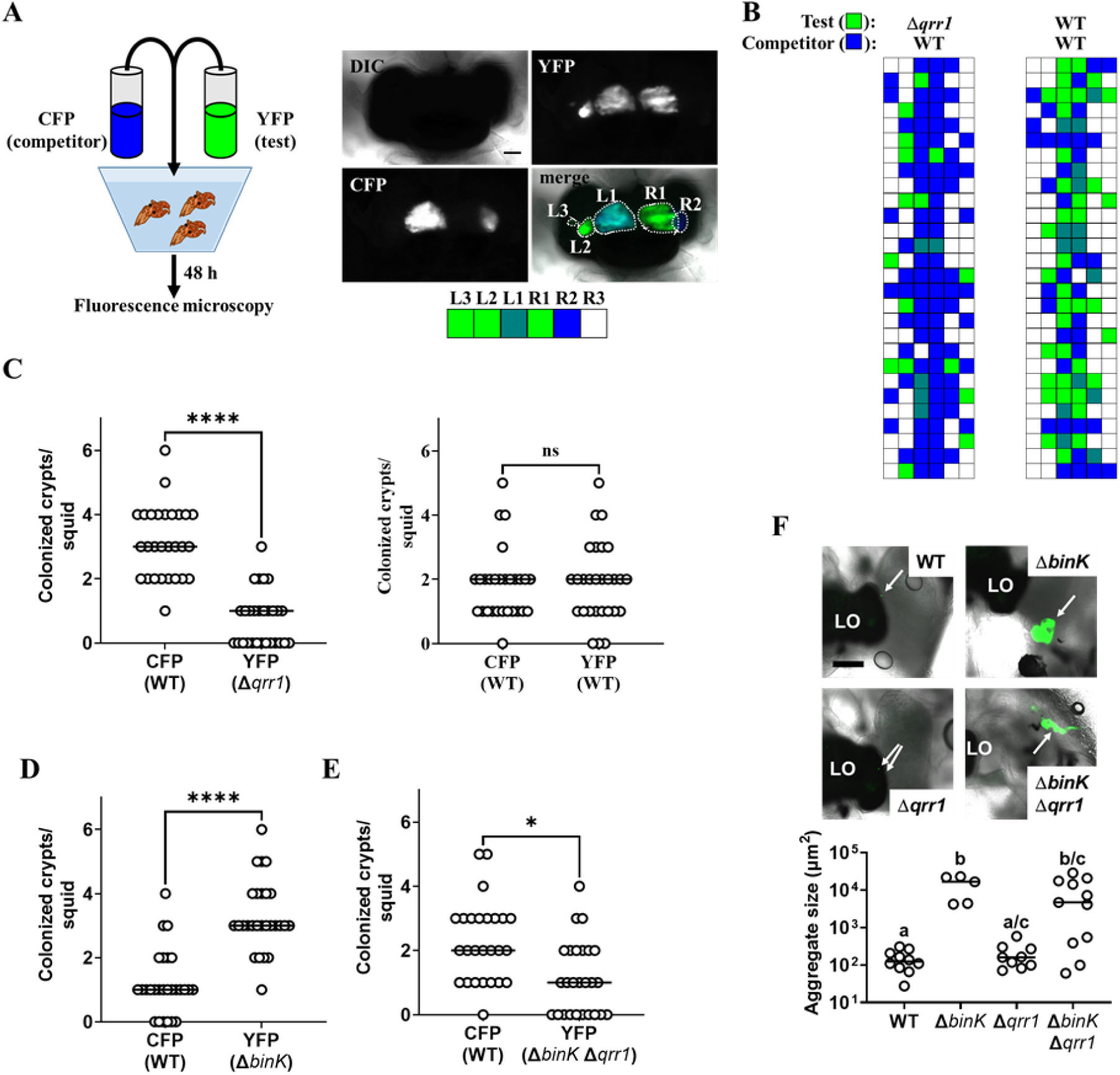
Qrr1 enhances the ability of *V. fischeri* to access crypt spaces. **(A)** *Left*, experimental design of squid co-inoculation assays with YFP-labeled test strain and CFP-labeled wild-type competitor strain. *Right*, example image montage illustrating a light organ featuring populations comprised of cells expressing YFP or CFP. Dotted line = boundary of an individual population. Scale bar = 100 µm. Boxes below the image indicate the strain type(s) present within each predicted crypt space of the ; blue = CFP^+^ YFP^-^, green = YFP^+^ CFP^-^, hatched = CFP^+^ YFP^+^, white = CFP^-^ YFP^-^. For panels B-E, experimental trials = 2. **(B)** *Left*, TIM305 (Δ*qrr1*) as test strain. *Right*, ES114 (WT) as test strain. N = 28. **(C)** Number of crypts colonized by indicated strains per squid in panel B. Wilcoxon test (**** = *p* < 0.0001, n.s. = not significant). **(D)** Δ*binK*. Number of crypts colonized by MJM2251 (Δ*binK*) as test strain. N = 27. Wilcoxon test (**** = *p* < 0.0001). **(E)** Δ*binK* Δ*qrr1*. Number of crypts colonized by EDR010 (Δ*binK* Δ*qrr1*) as test strain. N = 26. Wilcoxon test (* = *p* < 0.05). **(F)** Aggregation assay with ES114 (WT), MJM2251 (Δ*binK*), and EDR010 (Δ*binK* Δ*qrr1*) labeled with YFP. *Left*, merged brightfield and yellow fluorescence (green) images of aggregates (arrows) formed by indicated strains. LO = light organ. Scale bar = 200 µm. *Right*, quantification of aggregate size. Kruskal-Wallis (H = 16.79, *d.f.* = 3, *p* = 0.0008); Dunn’s *post-hoc* test with *p*-values corrected for multiple comparisons (same letter = not significant, a/c & b/c = *p* < 0.05, a/b = *p* < 0.01). Experimental trials: 2.

We next used this microscopy-based approach to investigate the Δ*binK* mutant. Following co-inoculation with the wild-type competitor strain, the Δ*binK* mutant occupied most of the crypt spaces (Fig. 3D), which suggests that Δ*binK* cells founded more populations than competitor cells and explains the previous observation of higher relative abundance of Δ*binK* cells in squid co-inoculated with those strain types (21). To determine whether Qrr1 impacts this effect, we also examined light organs of animals exposed to an inoculum mixed evenly with Δ*binK* Δ*qrr1* mutant and the wild-type competitor. The Δ*binK* Δ*qrr1* mutant occupied a minority of crypt spaces (Fig. 3E), which suggests that Qrr1 also promotes the ability of the Δ*binK* mutant to access crypt spaces. Because the Δ*binK* mutant forms large aggregates, we also considered whether Qrr1 affects this process by determining the extent to which the Δ*binK* Δ*qrr1* mutant could form aggregates. As expected, Δ*binK* formed larger aggregates than WT cells (Fig. 3F), which highlights the inhibitory role of BinK on aggregation formation that was previously reported (21). Most of the aggregates formed by the Δ*binK* Δ*qrr1* mutant were also large (Fig. 3F), which suggests that the impact of Qrr1 on aggregation formation is minimal. Taken together, these data suggest that the Δ*qrr1* allele is epistatic to the Δ*binK* allele during symbiosis establishment, which provides evidence that Qrr1 affects the ability of *V. fischeri* to enter a crypt space after the aggregation phase.

### The bEBP SypG activates σ^54^-dependent transcription of *P_qrr1_* in *V. fischeri*

To determine how *P_qrr1_* is activated in the Δ*binK* mutant, we considered factors known to promote transcription of *qrr1*. As with the *qrr* genes in other *Vibrionaceae* members (27), the promoter region of *qrr1* in *V. fischeri* (Fig. 4A) features nucleotides corresponding to the canonical -24 and -12 sites (TGGCA-N7-TGC) that facilitate binding by the alternative sigma factor σ^54^ (13). To test whether the *P_qrr1_* activity observed in the Δ*binK* mutant depends on σ^54^, we knocked out *rpoN*, which encodes σ^54^, from the Δ*binK* mutant and assessed *P_qrr1_*-*gfp* activity in the resulting Δ*rpoN* Δ*binK* double mutant. GFP levels in the double mutant were attenuated and comparable to the low levels of the Δ*rpoN* single mutant (Fig. 4B), which indicates that the activity of *P_qrr1_* of Δ*binK* cells depends on σ^54^.

**Figure 4:**
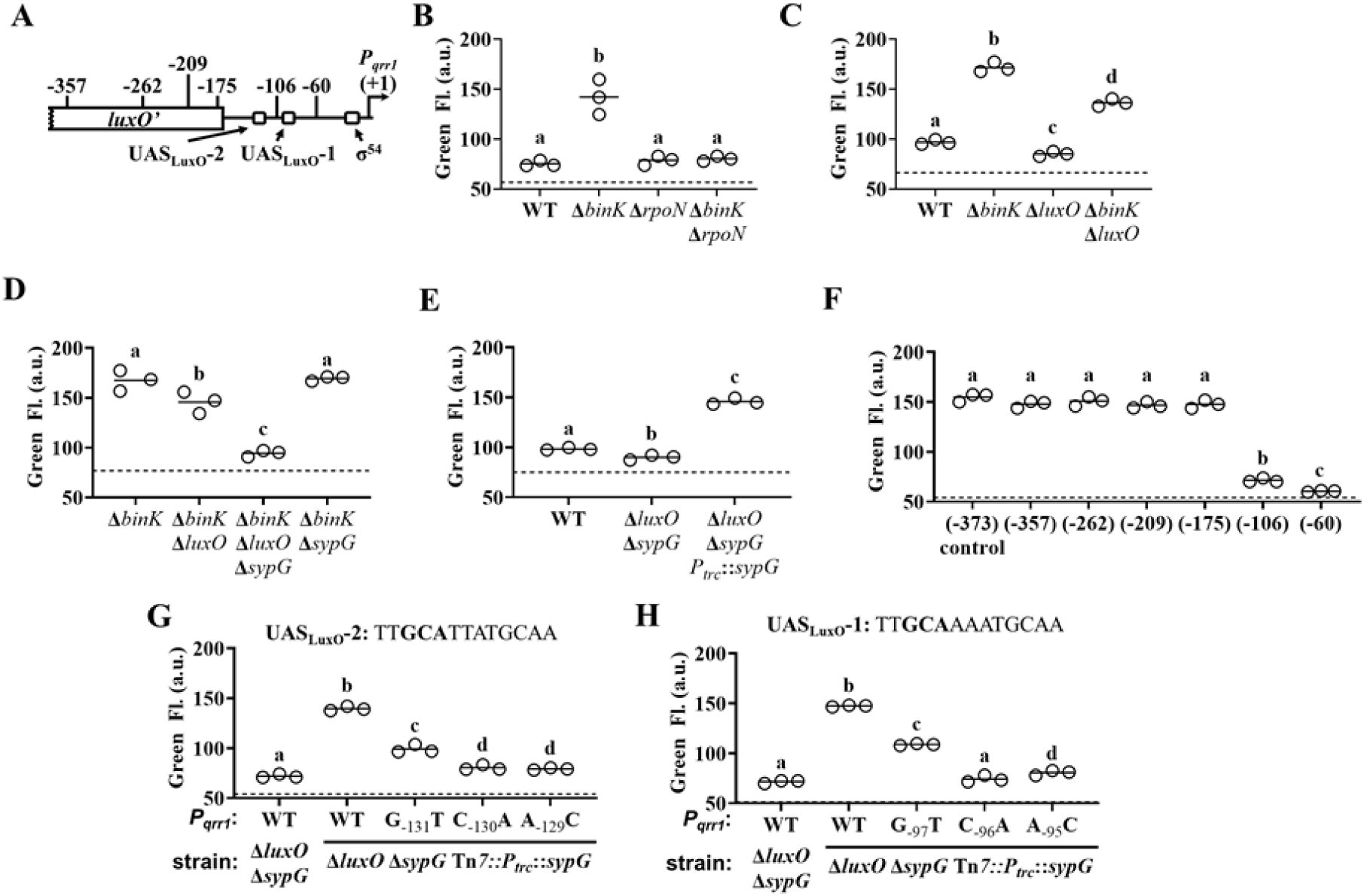
SypG activates σ^54^-dependent transcription of *qrr1*. **(A)** Elements within *Pqrr1* that facilitate σ^54^-dependent transcriptional activation. **(B)** Green fluorescence levels of ES114 (WT), MJM2251 (Δ*binK*), KRG004 (Δ*rpoN*), and KRG011 (Δ*binK* Δ*rpoN*) harboring pTM268 (*Pqrr1*::*gfp*). N = 3. Dotted line = autofluorescence cutoff. One-way ANOVA (F3,8 = 35.69, *p* < 0.0001); Tukey’s *post-hoc* test with *p*-values corrected for multiple comparisons (same letter = not significant, different letters = *p* < 0.001). **(C)** Green fluorescence levels of ES114 (WT), MJM2251 (Δ*binK*), TIM306 (Δ*luxO*), and (Δ*binK* Δ*luxO*) harboring pTM268 (*Pqrr1*::*gfp*). N = 3. Dotted line = autofluorescence cutoff. One-way ANOVA (F3,8 = 367.4, *p* < 0.0001). **(D)** Green fluorescence levels of MJM2251 (Δ*binK*), EDR009 (Δ*binK* Δ*luxO*), EDR014 (Δ*binK* Δ*sypG*), and EDR013 (Δ*binK* Δ*luxO* Δ*sypG*) harboring pTM268 (*Pqrr1*::*gfp*). N = 3. Dotted line = autofluorescence cutoff. One-way ANOVA (F3,8 = 60.66, *p* < 0.0001). **(E)** Green fluorescence levels of TIM313 (WT), EDS008 (Δ*luxO* Δ*sypG*), and EDS010 (Δ*luxO* Δ*sypG Ptrc*::*sypG*) harboring pEDR003 (*Pqrr1*::*gfp*) and grown on 150 µM IPTG. N = 3. Dotted line = autofluorescence cutoff. One-way ANOVA (F2,6 = 438.8, *p* < 0.0001). **(F)** Green fluorescence levels of EDS010 (Δ*luxO* Δ*sypG* Tn*7*::[*Ptrc*::*sypG erm*]) harboring *Pqrr1*::*gfp* reporter plasmids pEDR003, pEDR011 (-357), pEDR010 (-262), pEDR012 (-209), pEDR006 (-175), pEDR009 (-106), and pEDR008 (-60). N = 3. Dotted line = autofluorescence cutoff. One-way ANOVA (F6,14 = 411.7, *p* < 0.0001). **(G)** Green fluorescence levels of EDS008 (Δ*luxO* Δ*sypG*) and EDS010 (Δ*luxO* Δ*sypG* Tn*7*::*Ptrc*::*sypG*) harboring *Pqrr1*::*gfp* reporter plasmids pEDR003 (WT), pEDS004 (G-131T), pEDS005 (C-130A), and pEDS006 (A-129C). N = 3. Sequence corresponds to 13-bp UASLuxO-2, with nucleotides (-131)-(-129) that were individually mutated by site-directed mutagenesis shown in bold. Dotted line = autofluorescence cutoff. One-way ANOVA (F4,10 = 712.5, *p* < 0.0001). Experimental trials: 2. **(H)** Green fluorescence levels of EDS008 (Δ*luxO* Δ*sypG*) and EDS010 (Δ*luxO* Δ*sypG* Tn*7*::*Ptrc*::*sypG*) harboring *Pqrr1*::*gfp* reporter plasmids pEDR003, pEDS007 (G-97T), pEDS008 (C-96A), and pEDS009 (A-95C). N = 3. Sequence corresponds to 13-bp UASLuxO-1, with nucleotides (-97)-(-95) that were individually mutated by site-directed mutagenesis shown in bold. Dotted line = autofluorescence cutoff. One-way ANOVA (F4,10 = 335.3, *p* < 0.0001). Experimental trials: 2.

Transcriptional activation of σ^54^-dependent promoters critically depends on a bacterial enhancer binding protein (bEBP) interacting with nucleotides upstream of the promoter and hydrolyzing ATP to induce the conformation changes in the σ^54^-RNA polymerase complex that facilitate transcriptional activation (13). Therefore, we next considered whether the *P_qrr1_* activity observed in the Δ*binK* mutant depends on LuxO, which is the only bEBP known to activate σ^54^-dependent transcription of *P_qrr1_* (10).

While the GFP fluorescence level of a Δ*luxO* Δ*binK* mutant was lower than that of the Δ*binK* mutant (Fig. 4C), it was consistently higher than that of the wild-type strain, suggesting that LuxO is only partially responsible for σ^54^-dependent *P_qrr1_* activity in the Δ*binK* mutant.

The partial effect of LuxO described above suggests that a different bEBP also facilitates the σ^54^-dependent *P_qrr1_* activity observed in the Δ*binK* mutant. In addition to LuxO, the genome of ES114 encodes five other class I bEBPs: SypG, NtrC, VF_1401, FlrC, and VpsR. Of these other bEBPs, SypG stood out as a candidate for LuxO-independent activation of *P_qrr1_* for three reasons: i) SypG-dependent genes are elevated in the Δ*binK* mutant (21), ii) the primary structure of SypG is most identical to that of LuxO (Fig. S1) and predicted to form many of the structural features underlying LuxO function (15) (Fig. S2), and iii) WT cells harboring a multi-copy plasmid containing *sypG* exhibit elevated *P_qrr1_* activity (20). To test whether SypG affects the LuxO-independent *P_qrr1_* activity of Δ*binK* mutant cells, we constructed a Δ*binK* Δ*luxO* Δ*sypG* triple mutant. GFP fluorescence was lower in the triple mutant relative to the Δ*binK* Δ*luxO* mutant (Fig. 4D), which suggests that SypG promotes LuxO-independent *P_qrr1_* activity in cells lacking *binK*. Notably, *P_qrr1_* activity remained high in a Δ*binK* Δ*sypG* double mutant (Fig. 4D), which suggests that LuxO is the primary activator of *P_qrr1_* in the Δ*binK* mutant. Therefore, to test whether SypG is sufficient to activate *P_qrr1_*, we cloned *sypG* downstream of the IPTG-inducible *trc* promoter (*P_trc_*::*sypG*) and introduced this construct into a Δ*luxO* Δ*sypG* mutant harboring the *P_qrr1_-gfp* reporter. Induction of *sypG* expression in the double mutant resulted in GFP fluorescence (Fig. 4E), which suggests that SypG can activate *P_qrr1_* even in cells that encode BinK.

To further investigate how SypG activates *P_qrr1_* transcription, we evaluated the nucleotide sequence used in the construction of the *P_qrr1_*::*gfp* reporter for upstream activator sequences (UASs). In other *Vibrionaceae* members, the UAS of LuxO (UAS_LuxO_) corresponds to the 13-bp motif TTGCAWWWTGCAA, with W=A/T (27, 28). In ES114, the *P_qrr1_* region features two UAS_LuxO_ motifs centered at -93 bp (UAS_LuxO_-1) and -127 bp (UAS_LuxO_-2) relative to the transcriptional start site of *qrr1* (Fig. 4A), and nucleotide substitutions within either motif attenuate LuxO-dependent activation of *P_qrr1_* (Fig. S3). To determine which regions of the *P_qrr1_* region are necessary for SypG-dependent activation of *P_qrr1_*, a set of *P_qrr1_*::*gfp* reporter constructs with 5’-end truncations of various lengths was evaluated with *sypG* expression induced by IPTG. Two regions, one between -175 bp and -106 bp and a second between -106 bp and -60 bp, both contribute to SypG-dependent activation (Fig. 4F). Because each region contains a UAS_LuxO_, we also evaluated GFP fluorescence in *sypG*-induced cells harboring the reporters containing nucleotide substitutions within these motifs. *P_qrr1_* activity was attenuated for substitutions in either UAS_LuxO_ (Figs. 4G-H), which suggests that nucleotides corresponding to a UAS_LuxO_ also contribute to a UAS of SypG (UAS_SypG_). Consistent with hypothesis, each sequence exhibits similarity to the TTCTCANNNTGMDWN motif previously reported for SypG (29). Notably, the promoters of *sypA* and *sypP*, which are SypG-dependent genes in the *syp* locus, both show elevated activity in the Δ*binK* mutant, with only SypG contributing to the corresponding signals (Fig. S4). Furthermore, both *syp* promoters remained inactive in response to a hyperactive variant of LuxO (Fig. S4), which suggests that the ability of SypG and LuxO to activate transcription of a gene is specific to *P_qrr1_*. Taken together, these results suggest that SypG is a bEBP that can activate σ^54^-dependent transcription of Qrr1 in a manner independent of LuxO.

### Quorum sensing does not inhibit SypG-dependent activation of *qrr1*

Based on our finding that SypG activates transcription of *P_qrr1_*, we hypothesized that conditions that promote SypG activity would elevate the expression of Qrr1, which is significant because Qrr1- dependent regulation could occur under conditions of high cell density. To test this hypothesis, we first examined *P_qrr1_* activity in cells overexpressing RscS, which stimulates the expression of SypG-dependent genes (30). Using a plasmid containing the *rscS\** allele described above, RscS was overexpressed in *V. fischeri* strains engineered to encode a *P_qrr1_*::*gfp* reporter within its chromosome. When cell suspensions were spotted onto solid medium and incubated, the resulting surface structures featured pronounced heterogenous ridges (Fig. 5A), which comprise the wrinkled-colony phenotype that depends on expression of the *syp* locus (22). Green fluorescence was observed throughout the structure (Fig. 5A), particularly within the ridges, which suggests that *P_qrr1_* was activated from overexpressing RscS. In contrast, overexpression of RscS in a Δ*sypG* mutant resulted in smooth surface structures (Fig. 5A), which indicates the wrinkled-colony phenotype depends on a functional SypG, as previously reported (30). Furthermore, low green fluorescence was observed for the Δ*sypG* mutant (Fig. 5A), which indicates low *P_qrr1_* activity and suggests that SypG activation by RscS overexpression results in Qrr1 expression.

**Figure 5:**
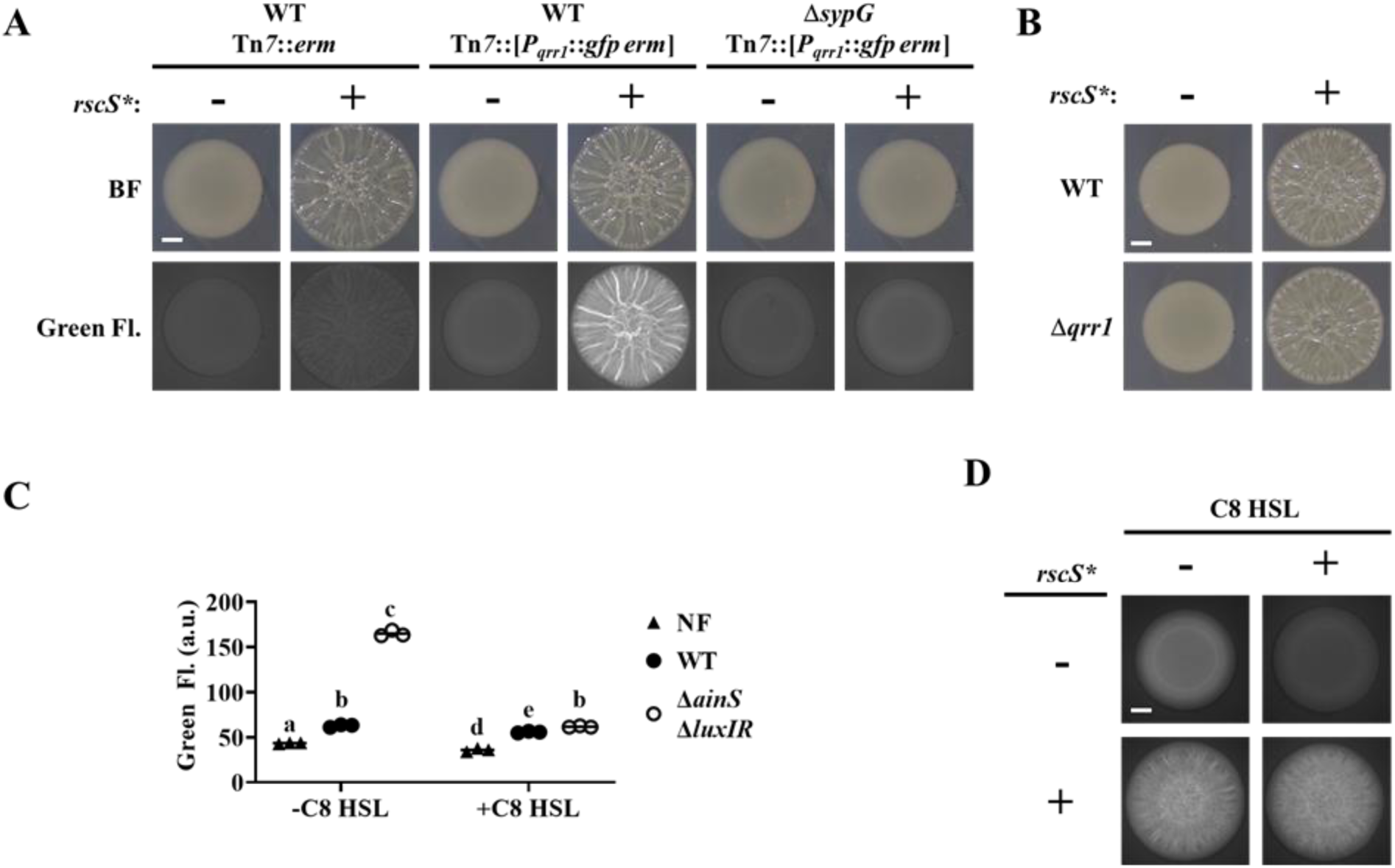
SypG activity overrides inhibition of *P_qrr1_* activity by quorum sensing. **(A)** Brightfield (top) and green fluorescence (bottom) images of representative spots of growth (N = 3) containing TIM313 (WT Tn*7*::*erm*), TIM303 (WT Tn*7*::[*P* ::*gfp erm*]), or EDS015 (Δ*sypG* Tn*7*::[*P* ::*gfp erm*]) harboring plasmid pKV69 (*rscS\** = -) or pKG11 (*rscS\** = +). Scale bar = 1 mm. **(B)** Brightfield images of representive spots of growth (N = 3) containing TIM303 (WT) or SSC (Δ*qrr1*) harboring plasmid pKG11 (*rscS\** = -) or pKG11 (*rscS\** = +). Scale bar = 1 mm. Experimental trials: 2. **(C)** Green fluorescence of ES114 (WT) and JHK007 (Δ*ainS* Δ*luxIR*) harboring *P* ::*gfp* reporter pTM268 (circles) and grown ± 100 nM C8 HSL (N = 3). ES114 harboring pVSV105 was used as a non-fluorescent control (NF). Two-way ANOVA revealed statistical significance for strain (F_2,12_ = 2809, p < 0.0001), C8 treatment (F_1,12_ = 2233, p < 0.0001), and their interaction (F2,12 = 1480, p < 0.0001); Tukey’s *post-hoc* test with *p*-values corrected for multiple comparisons (same letter = not significant, b/e = p < 0.05, a/d = *p* < 0.01, other combinations of different letters = *p* < 0.0001). Experimental trials: 2. **(D)** Green fluorescence images of representative spots of growth (N = 3) containing KRG016 (Δ*ainS* Δ*luxIR* Tn*7*::*P* ::*gfp*) harboring plasmid pKG11 (*rscS\** = +) or pKV69 (*rscS\** = -) on medium ± 100 nM C8 HSL. Scale bar = 1 mm. Experimental trials: 2.

However, a Δ*qrr1* mutant formed wrinkled colonies in response to overexpression of RscS (Fig. 5B), which suggests that Qrr1 does not promote the process of wrinkled colony formation.

We also investigated whether quorum sensing impacts SypG-dependent activation of *P_qrr1_*. In *V. fischeri*, signaling by the histidine kinase AinR in response to C8 HSL autoinducer results in lowered *P_qrr1_* activity (Fig. 1 and (11)). The low *P_qrr1_* activity observed in the spots of growth (Fig. 2) suggests that the level of C8 HSL is already elevated within the high cell density conditions, which would prevent our ability to detect a response to additional C8 HSL. Therefore, we introduced the *P_qrr1_*::*gfp* reporter into the chromosome of the Δ*ainS* mutant JHK007 (11), which does not produce the C8 HSL synthase AinS (31). In addition, JHK007 contains deletions of *luxI* or *luxR*, which contribute to an unknown mechanism that inhibits activation of *P_qrr1_* through AinR signaling (11). Consistent with this previous report, JHK007 showed elevated GFP fluorescence, which suggests high *P_qrr1_* activity in the absence of HSL-based autoinducers (Fig. 5C). Supplementing media with C8 HSL was sufficient to lower GFP fluorescence (Fig. 5C), which indicates that C8 HSL inhibits *P_qrr1_* activity. Using this experimental setup, we next assessed whether increased SypG activity could interfere with the ability of C8 HSL to inhibit *P_qrr1_* activity through the introduction of a plasmid harboring the *rscS\** allele. As expected, overexpression of RscS resulted in wrinkled colonies with elevated GFP fluorescence (Fig. 5D). However, the presence of C8 HSL did not alter the wrinkling phenotype and failed to lower GFP fluorescence, which suggests that *P_qrr1_* activity remained elevated in those spots. Taken together, these results suggest that SypG-dependent activation of *P_qrr1_* is insensitive to autoinducer and furthermore indicate a mechanism by which *V. fischeri* can express Qrr1 even when cells conduct quorum sensing.

### Diversity of SypG-dependent activation of *P_qrr1_* among *Vibrionaceae*

*V. fischeri* is a member of the Fischeri clade of *Vibrionaceae*, which includes five species that reside in seawater habitats as well as within squid and fish (32). The genomes of *Vibrionaceae* members commonly feature two chromosomes of unequal size, with the larger chromosome referred to as Chromosome 1 (33). The genomes of representative Fischeri taxa encode homologs of Qrr1 and LuxO on Chromosome 1 (Fig. S5 and Table 1) and SypG on Chromosome 2 (Table 1). Gene synteny associated with each locus across taxa suggests that the genes encoding Qrr1 and the bEBPs were passed vertically within the Fischeri lineage (Figs. S6A-B). For each taxon, alignment of the primary structures for the LuxO and SypG homologs revealed approximately 48% identity (Table 1 and Fig. S7). Among the five taxa, 44.4% (214/481) of residue positions in LuxO were identical to the corresponding SypG homolog (Figs. 6A-B and S8), which suggests that the functions associated with various domains of LuxO, including the regulatory linker and HTH domains, are also highly conserved in SypG. Together, these analyses based on bioinformatics suggest the possibility that SypG-dependent expression of Qrr sRNAs is conserved throughout the Fischeri clade.

**Figure 6:**
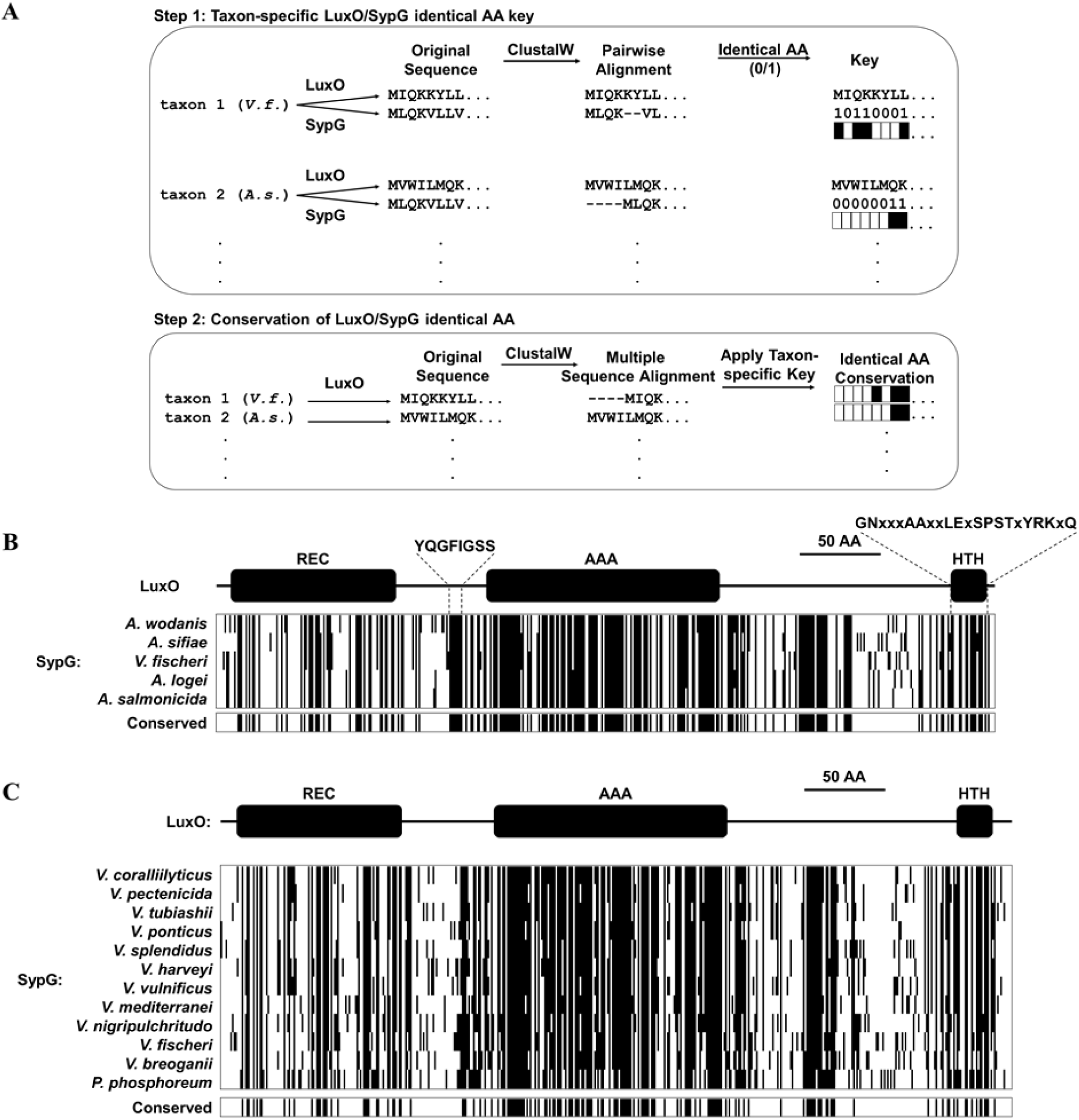
Conservation of SypG-LuxO structural features in the *Vibrionaceae* family. **(A)** Experimental design for visualizing the extent of identity conserved between the primary structures of LuxO and SypG homologs among different taxa. **(B)** Each block represents a multiple sequence alignment of LuxO homologs encoded within the indicated Fischeri clade members that has 481 amino acid positions including gaps. Positions marked by a black line indicate that the corresponding amino acid of the LuxO homolog is identical to that of SypG based on pairwise alignments. Shown below each block are the positions of amino acid identity that are conserved among the indicated taxa. **(C)** Each block represents a multiple sequence alignment of LuxO homologs encoded within the indicated *Vibrionaceae* members that has 489 amino acid positions including gaps. Positions marked by a black line indicate that the corresponding amino acid of the LuxO homolog is identical to that of SypG based on pairwise alignments. Shown below each block are the positions of amino acid identity that are conserved among the indicated taxa.

**Table 1:**
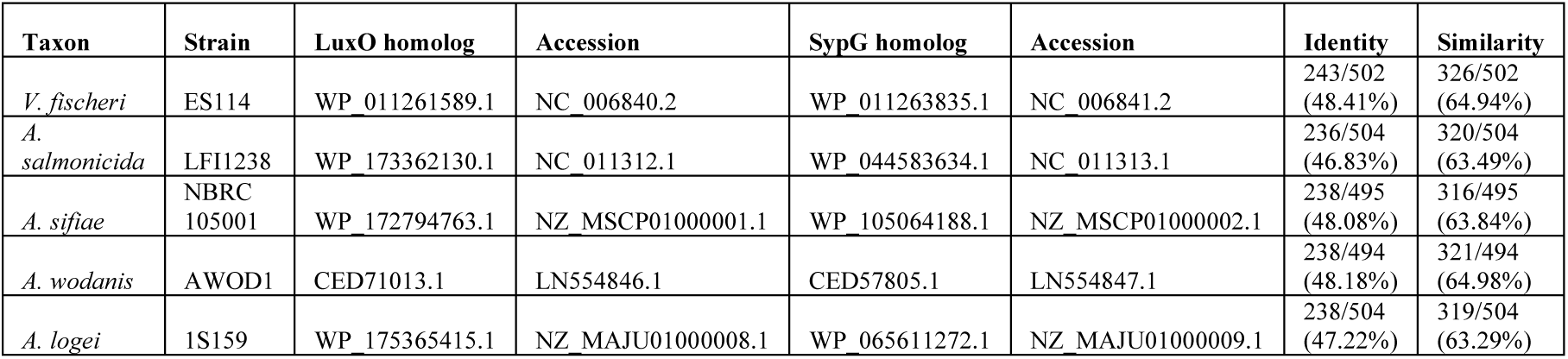
LuxO and SypG homologs in Fischeri clade.

We expanded our analysis to consider the *Vibrionaceae* family, which features species that are important in a variety of marine ecosystems, with many members able to cause disease in humans and other animals (34). Reconstruction of the evolutionary history of the *Vibrionaceae* family has resulted in 22 distinct clades, including Fischeri (32). All clades except Rumioensis feature taxa encoding a LuxO homolog (Table 2), with the corresponding *luxO* gene located on Chromosome 1 in the 17 taxa for which fully assembled genomes are available. Each of the remaining clades represented by the indicated taxa for which only contigs are available also featured a *luxO* gene, and gene synteny analysis of the neighboring genes suggests an arrangement consistent with its location on Chromosome 1 (Fig. S9). In addition, a putative Qrr is also encoded in opposite orientation of *luxO* in 20 of the 21 representative taxa that encode a LuxO homolog (Fig. S10), which suggests that the LuxO-Qrr regulatory system is highly conserved among *Vibrionaceae* members.

**Table 2:** LuxO and SypG homologs in *Vibrionaceae* clades.

| Clade | Taxon | LuxO homolog<sup>1</sup> | Accession | SypG homolog<sup>2</sup> | Accession | Identity<sup>3</sup> | Similarity<sup>3</sup> |
| --- | --- | --- | --- | --- | --- | --- | --- |
| Salinivibrio-Grimontia-Enterovibrio<sup>4</sup> | *G. hollisae* | WP_005503370.1 | NZ_CP014056 | N.D. | - | N.A. | N.A. |
| Rosenbergii | *P. lutimaris* | WP_107348500.1 | NZ_SNZO01000002 | N.D. | - | N.A. | N.A. |
| Profundum | *P. profundum* | WP_065814467.1 | NC_006370 | N.D. | - | N.A. | N.A. |
| Damselae | *Photobacterium damsela* subsp. *piscicida* | WP_086957069.1 | NZ_AP018045 | N.D. | - | N.A. | N.A. |
| Phosphoreum | *P. phosphoreum* | WP_045027808.1 | NZ_MSCQ01000001 | WP_105026695.1 | NZ_MSCQ01000001 | 237/499 (47.49%) | 310/499 (62.93%) |
| Fischeri | *V. fischeri* | WP_011261589.1 | NC_006840 | WP_011263835.1 | NC_006841 | 243/502 (48.41%) | 326/502 (64.94%) |
| Anguillarum | *V. anguillarum* | WP_026028983.1 | NC_022223 | N.D. | - | N.A. | N.A. |
| Rumoiensis | *V. rumoiensis* | N.D. | - | N.D. | - | N.A. | N.A. |
| Vulnificus | *V. vulnificus* | WP_011149911.1 | NC_014965 | WP_013571858.1 | NC_014965 | 247/508 (48.62%) | 320/508 (62.99%) |
| Diazotrophicus | *V. diazotrophicus* | WP_042486207.1 | NZ_POSL01000002 | N.D. | - | N.A. | N.A. |
| Gazogenes | *V. gazogenes* | WP_021019492.1 | NZ_CP018835 | N.D. | - | N.A. | N.A. |
| Porteresiae | *V. tritonius* | WP_068714228.1 | NZ_AP014635 | N.D. | - | N.A. | N.A. |
| Cholerae | *V. cholerae* | WP_001888250.1 | NC_002505 | N.D. | - | N.A. | N.A. |
| Haliotocoli | *V. breoganii* | WP_065209630.1 | NZ_CP016177 | WP_065210697.1 | NZ_CP016178 | 228/513 (44.44%) | 298/513 (57.70%) |
| Splendidus | *V. splendidus* | WP_004734031.1 | NZ_CP031055 | WP_065205220.1 | NZ_CP031055 | 237/511 (46.38%) | 314/511 (61.45%) |
| Pectenocida | *V. pectenocida* | WP_125320437.1 | NZ_RSFA01000020 | WP_125322971.1 | NZ_RSFA01000107 | 237/501 (47.31%) | 321/501 (64.07%) |
| Scophthalmi | *V. ponticus* | WP_075650093.1 | NZ_AP019657 | WP_075649540.1 | NZ_AP019657 | 229/506 (45.26%) | 319/506 (63.04%) |
| Nereis | *V. nereis* | WP_061781622.1 | NZ_BCUD01000001 | N.D. | - | N.A. | N.A. |
| Orientalis | *V. tubiashii* | WP_038550519.1 | NZ_CP009354 | WP_004748949.1 | NZ_CP009354 | 242/497 (48.69%) | 319/497 (64.19%) |
| Coralliilyticus | *V. coralliilyticus* | WP_019275536.1 | NZ_CP048693 | WP_021455926.1 | NZ_CP048693 | 242/503 (48.11%) | 322/503 (64.02%) |
| Harveyi | *V. harveyi* | WP_005444697.1 | NZ_CP009467 | WP_050907635.1 | NZ_CP009467 | 244/522 (46.74%) | 320/522 (61.30%) |
| Nigripulchritudo | *V. nigripulchritudo* | WP_022603175.1 | NC_022528 | WP_022550524.1 | NC_022528 | 247/508 (48.62%) | 331/508 (65.16%) |
| Mediterranei | *V. mediterranei* | WP_062462808.1 | NZ_CP018308 | WP_088875891.1 | NZ_CP018308 | 236/503 (46.92%) | 318/503 (63.22%) |

<sup>1</sup> N.D. (not detected) indicates that the top hit from BLAST was a bEBP other than LuxO.
<sup>2</sup> N.D. (not detected) indicates that the top hit from BLAST was a bEBP other than SypG.
<sup>3</sup> N.A. (not applicable) due to SypG homolog not detected.
<sup>4</sup> The *Salinivibrio*-*Grimontia*-*Enterovibrio* group is ancestrally related to the *Vibrionaceae* family and is included as an outgroup in this analysis.

Among the 21 taxa that encode a LuxO homolog, 12 of them also encode a SypG homolog (Table 2), and the corresponding *sypG* gene resides within a gene cluster that resembles the *syp* locus of *V. fischeri*. However, in contrast to the Fischeri clade, most taxa of other SypG-positive clades within the *Vibrionaceae* that have complete genomes encode the *syp* locus on Chromosome 1 (Table 2), which suggests the possibility that the *syp* locus was acquired by a progenitor of the Fischeri clade that arose after diversification from other *Vibrionaceae* lineages. Despite this possibility of independent acquisition events, the SypG homologs encoded by non-Fischeri taxa also exhibit high amino acid sequence identity to the corresponding LuxO homologues (Table 2), including the same structural features involved in regulating activity (Fig. 6C). To gain insight into the evolutionary history associated with the SypG homolog encoded by the Chromosome I of these other taxa, we evaluated the genomic context of *pepN*, which is genetically linked to the *syp* locus in several taxa but also highly conserved among all *Vibrionaceae*. Gene synteny analysis of *pepN* suggests that genome rearrangement likely contributed to certain taxa losing the *syp* locus, and consequently *sypG* (Fig. S11). Taken together, these observations suggest that while the *Vibrionaceae* lineage has undergone significant diversification with SypG, those taxa that encode both SypG and LuxO have the potential for SypG-dependent activation of Qrr sRNAs.

Finally, to test the possibility of SypG-dependent activation of *P_qrr1_* in taxa other than *V. fischeri*, we considered the fish pathogen *Aliivibrio salmonicida* strain LFI1238, which encodes a SypG homolog (SypG_As_) with nearly 47% identity to its LuxO homolog (Table 1 and Fig. S7).

Similar to *V. fischeri*, the genome of LFI1238 also features a single *qrr* gene (*qrr1_AS_*) with a promoter region (*P_qrr1AS_*) that contains motifs associated with σ^54^ binding and two UAS_LuxO_ sites (Fig. S12). The *sypG_AS_* gene was cloned downstream of *P_trc_* and ectopically expressed in the Δ*luxO* Δ*sypG* mutant of *V. fischeri*. Using a GFP reporter for the promoter of *qrr1_AS_* (*P_qrr1AS_*), we found that induction of *sypG_AS_* expression led to increased GFP fluorescence (Fig. 7), which suggests that SypG_AS_ can activate transcription of *P_qrr1AS_* and provides support that SypG-dependent expression of Qrr sRNAs can occur in other taxa within the *Vibrionaceae* family.

**Figure 7:**
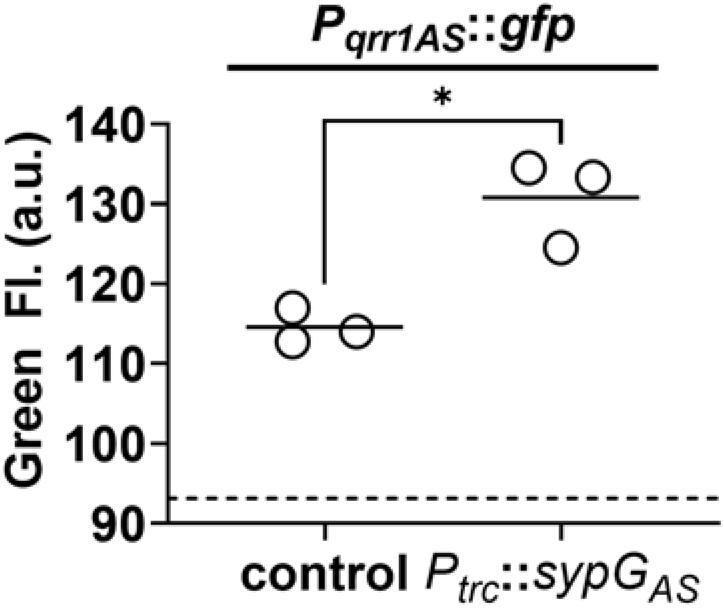
SypG-dependent transcription of *qrr1* for *A. salmonicida.* Green fluorescence of EDS008 (control) and EDS021 (Tn*7*::*sypG*_AS_) harboring pAGC003 (*P_qrr1AS_*::*gfp*). N = 3. Genotypes of both strains include Δ*luxO* and Δ*sypG* alleles, as well as *erm* integrated at the Tn*7* site. EDS008 harboring pVSV105 was used as a non-fluorescent control (dotted line). A paired *t*-test revealed significance between groups (* = *p* = 0.0325). Experimental trials: 2.

## Discussion

Quorum sensing enables all cells within a bacterial population to collectively express traits (7, 8). The traits regulated by quorum sensing are often energetically costly, and bacteria have adapted to inhibit their expression under non-quorum conditions. In *Vibrionaceae*, inhibitory factors include Qrr sRNAs, which post-transcriptionally repress expression of a transcription factor that promotes the cellular responses to quorum sensing (8). In this study, we discovered that *V. fischeri* has the potential to express Qrr1 even when responding to high concentrations of autoinducer (Fig 8A). Specifically, the bEBP SypG activates σ^54^-dependent transcription of *qrr1* in a manner that is independent of its primary bEBP LuxO. Transcriptional activation of *P_qrr1_* by SypG utilizes two UASs that overlap those sequences associated with LuxO-dependent activation. Together these findings reveal that *V. fischeri* has evolved to activate Qrr1 expression by either LuxO or SypG. The ability of SypG to activate σ^54^-dependent transcription of *P_qrr1_* in the presence of high autoinducer levels is significant because this regulatory link enables *V. fischeri* to bypass quorum sensing as a way to modulate the traits regulated by Qrr1.

**Figure 8:**
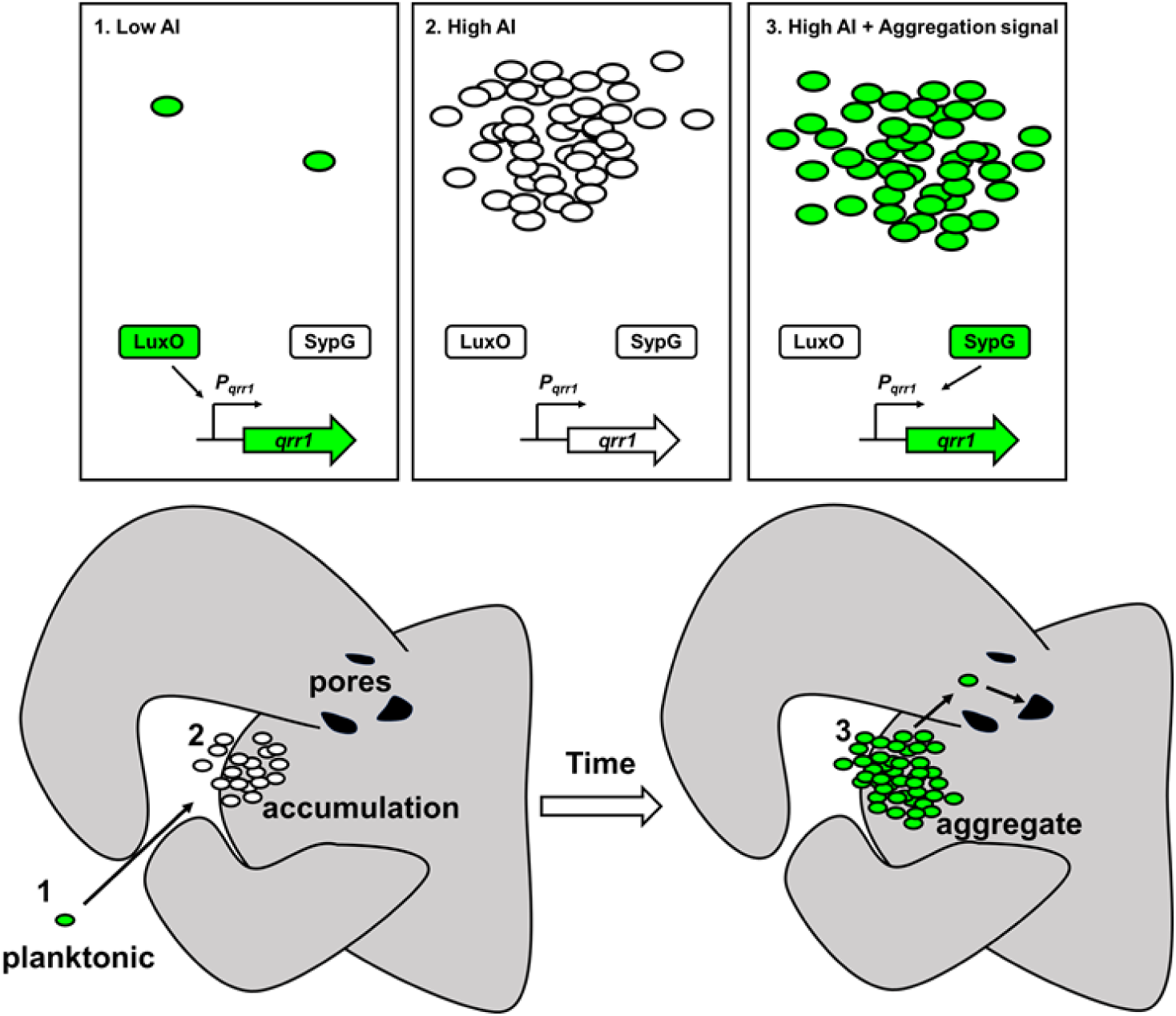
Model of dual bEBP control over Qrr1 expression in *V. fischeri*. **(A)** *Left,* cells in an environment with low autoinducer concentration, *e.g.*, low cell density, will express Qrr1 by activating LuxO through the quorum-sensing pathway. *Middle*, cells in an environment with high autoinducer concentration, *e.g.*, high cell density, will have low Qrr1 levels due to the inactive state of LuxO. *Right*, even under conditions of high autoinducer concentration, expression of Qrr1 can occur if SypG is activated by the aggregation pathway. **(B)** Model of initial entry of *V. fischeri* into the light organ. Planktonic cells within the environment express Qrr1 due to low autoinducer levels (panel A, box 1). Motion of the cilia associated with the appendages sweep bacteria into a stagnant zone, where they locally accumulate (panel A, box 2), which has the potential to lower Qrr1 expression. Within a few hours, the cells have formed aggregates that depend on SypG (panel A, box 3), which induces Qrr1 expression to prime cells for entry into the pores.

When during the life cycle of *V. fischeri* would SypG-dependent activation of *P_qrr1_* be important? SypG activates σ^54^-dependent transcription of the *syp* locus, which enables *V. fischeri* to secrete polysaccharides that form an extracellular matrix (30). Production of extracellular polysaccharides is necessary for *V. fischeri* to form the cellular aggregates on the surface of the light organ while initiating symbiosis (22, 29, 35). In culture, *syp*-dependent biofilm formation, which has been used to model the aggregation observed *in vivo*, depends on SypG activating expression of all four operons within the *syp* locus (29). While this study implicates *qrr1* as a member of the SypG regulon, its activation does not appear to contribute to biofilm formation (Fig. 5B), which suggests that the Δ*qrr1* mutant can form aggregates prior to establishing symbiosis, in contrast to mutants containing deletions in other SypG- dependent genes. However, a Δ*qrr1* mutant shows fewer crypt populations relative to wild-type cells (Fig. 3), which suggests that regulation by Qrr1 enhances the ability of *V. fischeri* to access a crypt space. Taken together, these findings support a model by which the environmental *V. fischeri* cells collected by the squid host express Qrr1 via SypG while forming *syp*-dependent aggregates along the light organ surface (Fig. 8B). SypG-dependent activation of Qrr1 would prime cells to express certain traits that are enhanced by this sRNA, *e.g.*, cellular motility, precisely when the transition from the aggregate stage to light organ entry occurs. Notably, the ability of SypG to activate transcription of *P_qrr1_* makes the phosphorylation state of LuxO, and by extension, the corresponding quorum-sensing signaling pathway, irrelevant for expressing Qrr1 during this critical stage of initiating symbiosis. Interestingly, previous work has demonstrated that Qrr1 can be expressed under conditions of high cell density through the overexpression of SypK (20), which is a putative oligosaccharide encoded by the *syp* locus. The current model is that SypK, which is predicted to localize to the inner membrane, activates *P_qrr1_* transcription by stimulating the LuxP/Q complex to trigger LuxO activity. More recently, it was also shown that a small molecule produced within RscS-induced wrinkled colonies promotes bioluminescence production (36), which suggests *V. fischeri* may feature additional connections between aggregation and quorum-sensing pathways. The finding that SypG can also activate *P_qrr1_* further expands the hypothesis that conditions that activate the *syp* locus, *e.g.*, when *V. fischeri* is initiating symbiosis, lead to the expression of Qrr1 as a mechanism to prime cells for host colonization.

Homologs of LuxO and Qrrs are encoded by most *Vibrionaceae* genomes, which underscores their biological significance in regulating the traits associated with quorum sensing. Over half of the *Vibrionaceae* clades feature taxa that also encode SypG homologs with a high degree of amino acid identity/similarity to the corresponding LuxO homologs (Table 2 and Fig. 6). Such high similarity among primary structures is likely to promote higher-order structures within SypG that function similarly to those of LuxO. For instance, the 1.6 Å-resolution crystal structure derived from a partial-length construct of *V. angustum* LuxO (PDB entry 5EP0) features a linker region between the REC and AAA+ domains that sterically occludes nucleotide binding thereby preventing the ATP hydrolysis necessary for remodeling the RNAP-σ^54^ complex to initiate transcription (15). A glycine conserved among all LuxO homologs both stabilizes this linker and occupies the active site, and, consistent with its predicted inhibitory role, substitution of the corresponding glycine in the LuxO homolog of *V. cholerae* with glutamate (G145E) results in increased LuxO activity (15). The analysis presented here shows that the primary structure of the linker is broadly conserved among the SypG homologs that are encoded by various *Vibrionaceae* members (Figs. 6). To our knowledge, SypG represents the only other bEBP aside from LuxO predicted to contain this regulatory linker. Notably, examination of other *Vibrionaceae* clades did reveal some intriguing exceptions, *e.g.*, the position corresponding to a glycine within the linker is an asparagine in *V. splendidus* (N141) and an aspartate in *V. mediterranei* (D141). Both substitutions involve residues that are larger than glycine, which is the only residue that can fit within the active site of the *V. angustum* LuxO structure (15). Therefore, the SypG homologs of *V. splendidus* and *V. mediterranei* are likely to exhibit constitutive activity or feature other adaptations that accommodate for the altered linker structure. Future crystallographic and biochemical studies of these SypG homologs are necessary to test these possibilities. In addition, investigation into how each SypG homolog affects various traits in the corresponding taxon will provide insight into the various ecological roles of the *syp* locus among the *Vibrionaceae* family.

Because bEBPs are critical for σ^54^-dependent transcriptional activation (14) and their activity is usually controlled by signal transduction networks that sense environmental stimuli (13), these specialized transcription factors also offer opportunities to engineer tightly controlled gene-regulatory modules for use in synthetic biology applications. For instance, the ability of LuxO and SypG to each activate transcription of *P_qrr1_* presented here resembles an OR logic gate that permits gene expression when either one or both of the bEBPs are active (*e.g.*, Fig. 4D). Molecular OR logic gates have been proposed as important components in engineering therapeutic bacteria that will deliver a drug when certain environmental conditions are satisfied (37). To our knowledge, evidence of different bEBPs activating the same gene has only been observed when the promoter exhibits distinct UASs that are specific for one or another bEBP. For instance, the σ^54^-dependent *dctA* gene of *Sinorhizobium meleloti* retains 20% transcriptional activity in a mutant lacking the *dctD* gene encoding the primary bEBP (38). This residual activity has been attributed to the bEBP NifA, for which potential UAS sites were identified within the *dctA* promoter region at sequences other than those associated with DctD binding (38, 39).

Despite the SypG and LuxO utilizing the overlapping UASs upstream of *P_qrr1_*, it appears that the σ^54^- dependent promoters of the *syp* locus can be activated by SypG but not by LuxO (Fig. S4). This discovery expands the utility of the LuxO and SypG for synthetic biology with the *syp* promoters being appropriate for controlling gene expression with SypG alone. Consequently, determining the mechanism by which the *syp* promoters are insulated from LuxO in *V. fischeri* will not only reveal molecular insight into symbiont biology but will also further expand the utility of bEBPs in synthetic biology applications.

Full understanding of the structure-function relationship underlying the OR logic gate described in this study will require further investigation into the molecular details by which LuxO and SypG activate *P_qrr1_*. For instance, determining how the HTH domain of each bEBP interacts with DNA will provide insight into whether competition between LuxO and SypG can affect dynamics of *P_qrr1_* activity under different environmental conditions. Furthermore, the high degree of identity within the AAA+ domains may facilitate the assembly of LuxO-SypG heterohexamers with activity levels that are different from their homohexameric forms. Because the activity of each bEBP is linked to distinct signal transduction systems, this finding expands understanding of the environmental conditions that impact the cellular physiology of *V. fischeri*.

### Methodology

#### Strains and plasmids

*V. fischeri* strains and plasmids used in this study are listed in Table 3. For cloning, *E. coli* strains Top10 and S17-1λpir were used. Primers used in the construction of strains and plasmids are listed in Table 4.

**Table 3:**
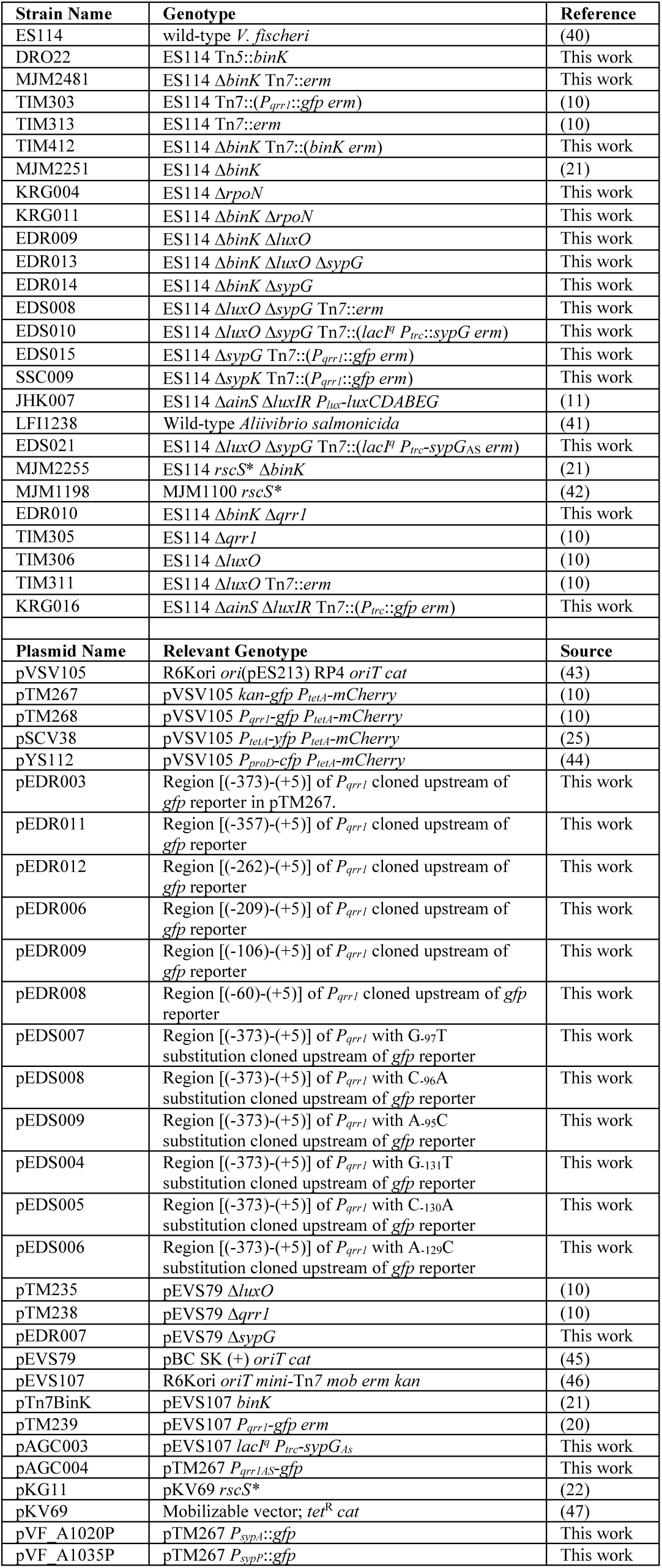
Strains and plasmids used in this study.

**Table 4:**
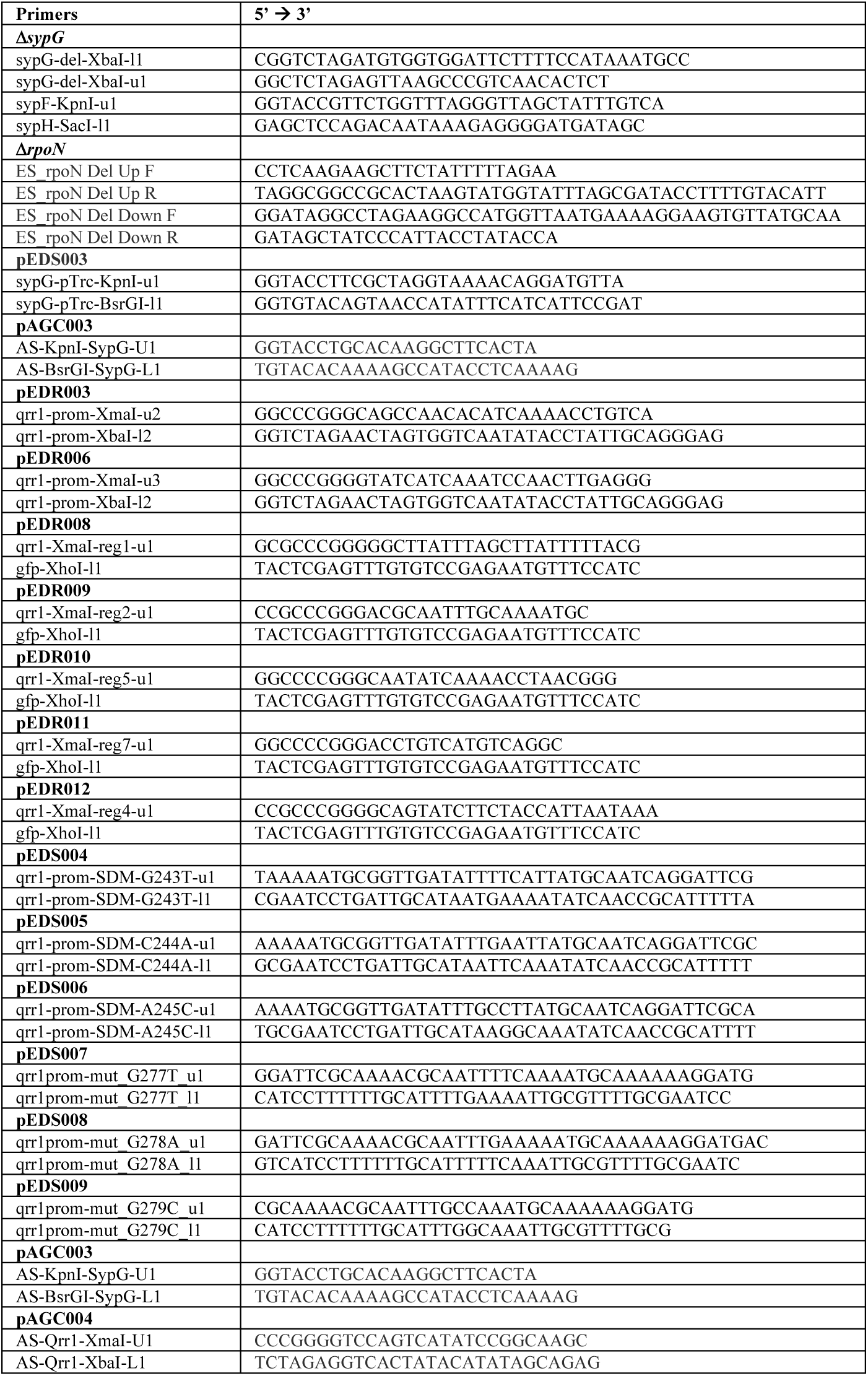
Primers used in this study.

#### Media and growth conditions

*V. fischeri* strains were grown at 28°C under aerobic conditions in LBS (Luria-Broth Salt) medium [1% (wt/vol) tryptone, 0.5% (wt/vol) yeast extract, 2% (wt/vol) NaCl, 50mM Tris-HCl (pH 7.5)] or SWT (seawater-tryptone) medium (48) with Instant Ocean (Blacksburg, VA) replacing seawater.

### Molecular biology

Construction of deletion alleles. Deletion alleles for *luxO*, *qrr1*, and *sypG* were introduced into strains by performing allelic exchange, as described previously (10). Construction details of plasmids pTM235 and pTM238 that feature Δ*luxO* and Δ*qrr1*, respectively, were described elsewhere (10). Plasmid pEDR007, which encodes the deletion allele of *sypG* (Δ*sypG*) that lacks the codons encoding residues 49-478, was constructed by first amplifying by PCR from ES114 genomic DNA regions of homology upstream (primers sypF-KpnI-u1 & sypG-del-XbaI-l1) and downstream (primers sypG-del-XbaI-u1 & sypH-SacI- l1) of *sypG* and then cloning them into pEVS79 via KpnI/SacI. The deletion allele for *rpoN* (Δ*rpoN*) features the entire *rpoN* gene (1,470 bp) replaced with the 78-bp FRT scar was introduced into strains by SOE PCR and recombineering mutagenesis (49) by generating regions of homology upstream (primers ES_rpoN Del Up F & R) and downstream (primers ES_rpoN Del Down F & R).

Chromosomal integration. Plasmids pEVS107, pTn7binK, pTM239, pEDS003, and pAGC003 were used to introduce genetic content in single copy into the chromosome at the Tn*7* site, as described elsewhere (46). Plasmid pEDS003 was constructed by first amplifying *sypG* by PCR from ES114 genomic DNA (primers sypG-pTrc-KpnI-u1 & -BsrGI-l1) and cloning the product downstream of the *P_trc_* promoter in pTM318 via KpnI/BsrGI. Plasmid pAGC003 was constructed in similar fashion using the amplicon (primers AS-KpnI-SypG-U1 & -BsrGI-SypG-L1) generated from LFI1238 genomic DNA.

Promoter transcriptional reporters. Plasmids pEDR003 and pEDR006 were constructed by amplifying the *P_qrr1_* region from ES114 genomic DNA by PCR (primers qrr1-prom-XmaI-u2 & -XbaI-l2 and -XmaI-u3 & XbaI-l2, respectively) and cloning the products upstream of *gfp* in pTM267 via XmaI/XbaI. Reporter plasmids pEDR011, pEDR010, pEDR012, pEDR009, and pEDR008, which contain truncated *P_qrr1_* regions, were constructed by amplifying from pEDR003 by PCR (reverse primer gfp-XhoI-l1 and respective forward primers qrr1-XmaI-reg7-u1, -reg5-u1, -reg4-u1, -reg2-u1, and -reg1-u1) and cloning the resulting products into pTM267 via XmaI/XhoI. Plasmid pAGC004, which contains the *P_qrr1AS_*-*gfp* reporter, was constructed by amplifying the Pqrr1AS region from LFI1238 genomic DNA by PCR (primers AS-Qrr1-XmaI-U1 & -XbaI-L1) and cloning the product into pTM267 via XmaI/XbaI.

Site-directed mutagenesis. The amplicon generated for pEDR003 (primers qrr1-prom-XmaI-u2 & XbaI- l2), which contains the *P_qrr1_* region, was cloned into pCR-blunt (ThermoFisher) and used as a template for site-directed mutagenesis. Primers listed for pEDS004, pEDS005, pEDS006, pEDS007, pEDS008, and pEDS009 were used to conduct PCR with Pfu Ultra (Agilent) for 18 cycles. The reaction was subjected to DpnI digest, transformed by electroporation into Top10 *E. coli* cells, and validated by sequencing before subcloning into pTM267 via XmaI/XbaI.

#### Promoter-activity spotting assays

Starter cultures of *V. fischeri* strains were grown overnight in LBS broth supplemented with 2.5 µg/mL chloramphenicol. For each culture, a 1-mL sample was prepared by adjusting its turbidity to an OD_600_ equivalent to 1.0. To initiate the assay, a 2.5-µL sample of the cell suspension was placed onto the surface of LBS agar supplemented with 2.5 µg/mL chloramphenicol (and 150 µM IPTG where indicated) and incubated at 28°C. After 24 h, the spots were examined at 4X magnification using an SZX16 fluorescence dissecting microscope (Olympus) equipped with an SDF PLFL 0.3X objective and both GFP and mCherry filter sets. Images of green fluorescence and red fluorescence of the spot were captured using an EOS Rebel T5 camera (Canon) with the RAW image format setting. Image analysis was performed using ImageJ, v. 1.52a (NIH) as follows. First, images were converted to RGB TIFF format using the DCRaw macro, with the following settings selected: use_temporary_directory, white_balance = [Camera white balance], do_not_automatically_brighten, output_colorspace = [sRGB], read_as = [8-bit], interpolation = [High-speed, low-quality bilinear], and half_size. For each spot, the green channel of the green fluorescence image was used for quantifying GFP fluorescence, and the red channel of the mCherry fluorescence image was used for quantifying mCherry fluorescence. The region of interest (ROI) corresponding to the spot was identified in the red channel by thresholding, and this ROI was used to determine the mean red and green fluorescence levels for each spot. A non-fluorescent sample (pVSV105/ES114) was used to determine the levels of cellular auto-fluorescence. A one-way ANOVA with Dunnett’s multiple comparisons test was performed to determine whether groups were significantly different than the non-fluorescent control group. The fold change in fluorescence between two groups was determined by first subtracting auto- fluorescence levels from each group mean and then calculating the ratio of the differences.

#### Bioluminescence assay

Starter LBS cultures of the indicated *V. fischeri* strains were grown overnight and then sub-cultured 1:100 into seawater tryptone (SWT) medium. At indicated time points, turbidity (OD_600_) and luminescence (RLUs) measurements were collected using a Biowave CO8000 Cell Density Meter and a Promega GloMax 20/20 luminometer, respectively. Specific luminescence for each sample was calculated by normalizing each luminescence measurement with the corresponding turbidity measurement.

#### Light-organ colonization assay

Starter cultures of the indicated *V. fischeri* strains were initiated with LBS medium supplemented with 2.5 µg/mL chloramphenicol for plasmid maintenance. Following overnight incubation, culture samples were normalized to an OD_600_ = 1.0 and diluted 1:100 in fresh medium. After cultures had reached OD_600_ = 1.0, they were diluted into filter-sterilized Instant Ocean seawater (FSSW). For each group, freshly hatched juvenile squid (*Euprymna scolopes*) derived from wild-caught adult animals collected in Oahu, HI and maintained in a mariculture facility were exposed collectively to an inoculum mixed evenly with cell suspensions of the indicated *V. fischeri* strains. The total cellular abundance and ratio of strain types in each inoculum were determined by plating serial dilutions and using a fluorescence dissecting microscope to count the resulting colonies exhibiting YFP and CFP fluorescence. Inoculum levels ranged between 4 x 10^3^ and 1 x 10^5^ CFU/mL and corresponding ratios were not significantly different from 1.0. After being exposed to the inoculum for 3.5 h, squid were washed three times in FSSW and then housed individually in vials containing 4 mL FSSW. Each day, squid were transferred to vials containing fresh FSSW. At 44 hours post-inoculation, squid of each group were combined and anesthetized on ice with 5% ethanol/FSSW and then fixed as a group in marine phosphate buffer containing 4% paraformaldehyde at 4°C. After 24 h, squid were washed four times with marine phosphate buffer and dissected to reveal the light organ. For each light organ, images of YFP, CFP, and DIC were acquired using a 780 NLO confocal microscope (Carl Zeiss AG, Jena, Germany) equipped with a 10X water lens and pinholes set to maximum to mimic epi-fluorescence conditions. The YFP and CFP fluorescence images of each light organ were visually examined in conjunction with the DIC image to score each region associated with a crypt space for fluorescence signal. Animal experiments were performed using protocol approved by the Institutional Animal Care and Use Committee at Penn State University (#PROTO202101789).

#### Aggregation assay

Starter LBS + 2.5 µg/mL chloramphenicol cultures of indicated strains harboring pSCV38 were diluted 1:100 into fresh medium and grown to an OD_600_ = 1.0. Cells were washed twice with each step consisting of centrifugation at 5,000 x *g* for 2 minutes, aspiration of the supernatant, and resuspension of the pellet into FSSW. The assay was initiated by exposing squid as a group to 5.0 x 10^6^ CFU/mL. After 3.5 hours, squid were anesthetized by placing on ice for 5 minutes and then exposing them to 3% ethanol/FSSW for at least 15 minutes. The light organ was exposed by dissection with forceps and imaged using fluorescence microscopy. Each light organ was scored for aggregates by assessing the green fluorescence image of each side for the presence of a particle. Aggregate size was determined using the default IsoData auto-threshold method of the threshold tool in ImageJ to generate a binary image from the green fluorescence image, which was then subjected to the analyze particles command, with pixel^2 size range set to 10-infinity, to measure the area of each particle.

#### Wrinkled colony assay

Starter cultures of *V. fischeri* strains harboring either pKG11 (*rscS\**) or pKV69 (vector) were grown overnight in LBS broth supplemented with 2.5 µg/mL chloramphenicol. For each culture, a 1-mL sample was prepared by adjusting its turbidity to an OD_600_ equivalent to 1.0. To initiate the assay, a 2.5-µL sample of the cell suspension was placed onto the surface of LBS agar supplemented with 2.5 µg/mL chloramphenicol and incubated at 25°C. After 24 h, the spots were examined at 4X magnification using an SZX16 fluorescence dissecting microscope (Olympus) equipped with an SDF PLFL 0.3X objective and either a GFP filter (green fluorescence) or no filter (brightfield). Images were acquired as described in the promoter-activity spotting assay above.

#### Statistical analysis

Except where indicated in the figure legend, experiments were performed at least three times. We define biological replicates as biologically distinct samples showing biological variation, and technical replicates as repeated measurements of a single sample. The number of biological replicates (N) is listed in figure legends. All statistical tests were performed in GraphPad Prism version 9.3.1 and listed in figure legends. Justification for statistical tests was determined by performing a Shapiro-Wilk test for normality on group data (or log-transformed data). Experiments in which normality failed (*p*-value ≥ 0.05) were statistically analysed using nonparametric statistical tests.

#### Gene synteny analysis

Analysis of gene synteny was performed by downloading GenBank files containing the indicated sequences from NCBI and subjecting them to the progressiveMauve algorithm, which identifies locally colinear blocks (LCBs) that are genomic segments that are conserved independent of rearrangements due to recombination. The following parameters were selected for each run: default seed weight, determine LCBs, full alignment, iterative refinement, and sum-of-pairs LCB scoring.

#### Protein alignments

Protein sequences were downloaded as FASTA format from NCBI and pasted directly into the Alignment Explorer tool of MEGA X (50). Alignments were performed using ClustalW, with Gap Opening Penalty = 10.00 and Gap Extension Penalty = 0.10 and 0.20 for pairwise and multiple sequence alignments, respectively. Alignments were exported in .fas format. For pairwise alignments, the identity and similarity values were determined using the Ident and Sim program of the Sequence Manipulation Suite (SMS) (50). Alignment displays were generated using the Color Align Conservation program of SMS, with similar amino acid groups defined as GAVLI, FYW, CM, ST, KRH, DENQ, P.

To visualize the positions of residues that are identical between LuxO and SypG homologs across a set of taxa, a multisequence alignment of the LuxO homologs encoded by those taxa was first generated. Each pairwise alignment was used to generate a key that indicates for each residue in LuxO whether the corresponding position within the alignment contains an amino acid that is identical (labeled as 1) or not identical (labeled as 0). The keys from the pairwise alignments were used to replace the amino acid letters within the LuxO multisequence alignment with the identical/not identical values. Using Excel, cells containing a 1 were formatted with black fill and those cells containing a 0 were formatted with white fill. The resulting table grid was used to generate the corresponding image shown in this report.

The consensus array was generated in similar fashion after determining which positions across rows within the alignment contained a value of 1.

## Acknowledgements

This work was supported by National Institute of General Medical Sciences Grants R01 GM129133 (to T.I.M.) and R35 GM119627 (to M.J.M.), Howard Hughes Medical Institute Gilliam Fellowship (to E.D.S. and T.I.M.), and National Institute of Allergy and Infectious Diseases Fellowship F32 AI 147543 (to K.R.G.).

## Material Availability Statement

Reasonable requests for plasmids and strains can be made to corresponding author (T.I.M.).

---

**Figure S1:**
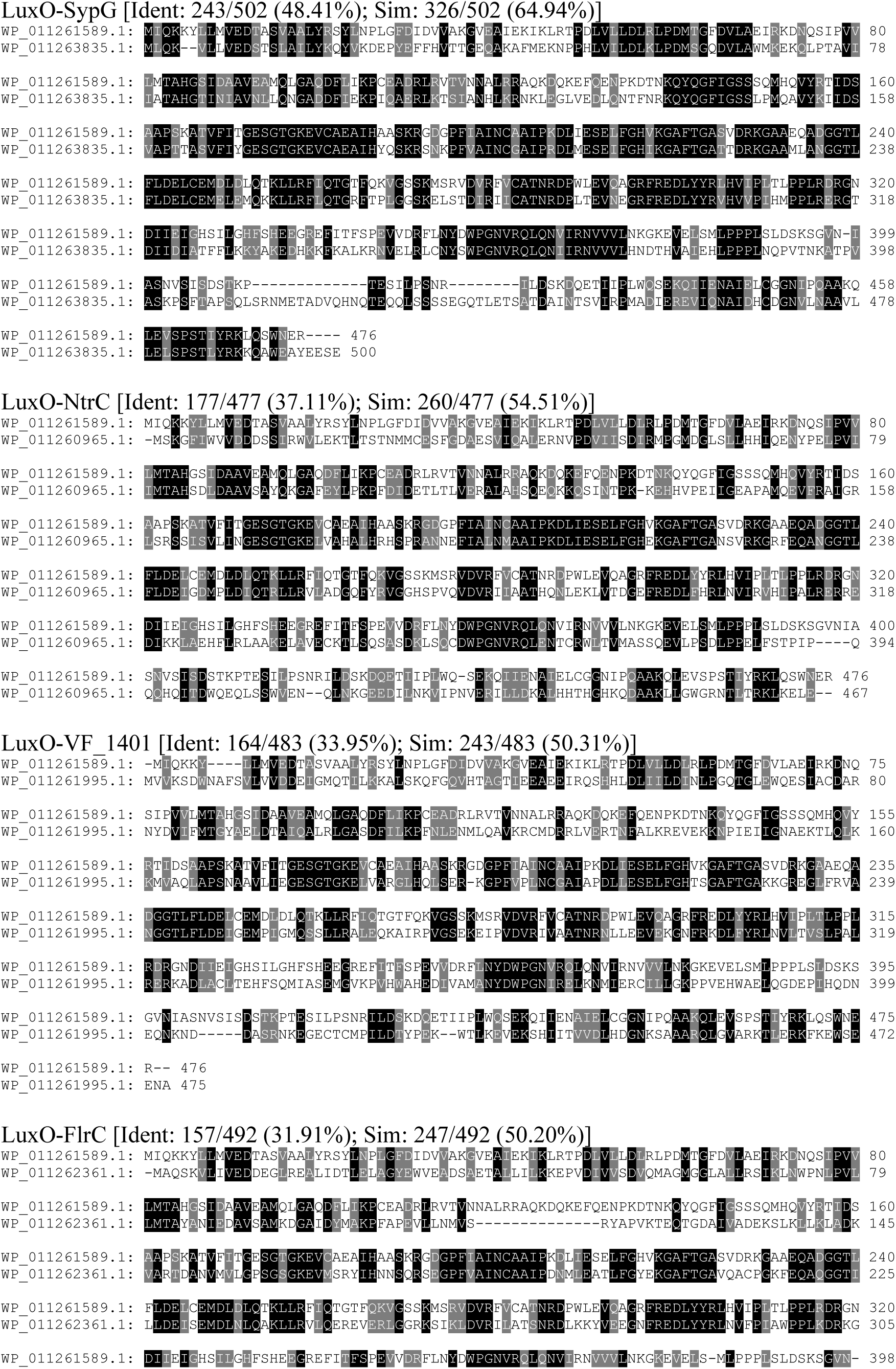
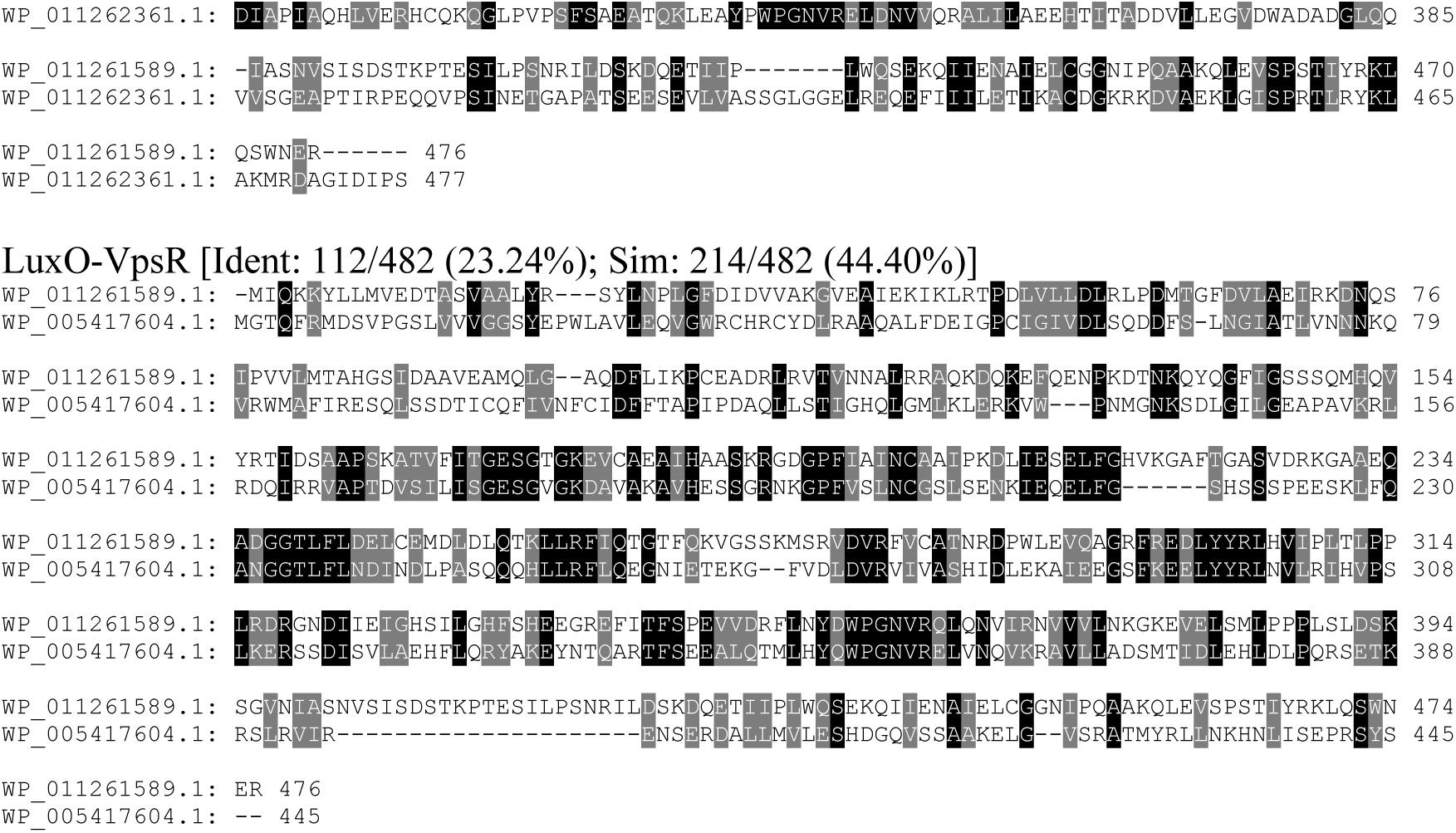
Comparisons of LuxO with other Class I bEBPs encoded by *V. fischeri*. Pairwise alignments of LuxO (top sequence) with indicated bEBP (bottom sequence) encoded by *V. fischeri* strain ES114. Amino acids highlighted in black or gray indicate identity and similarity, respectively.

**Figure S2:**
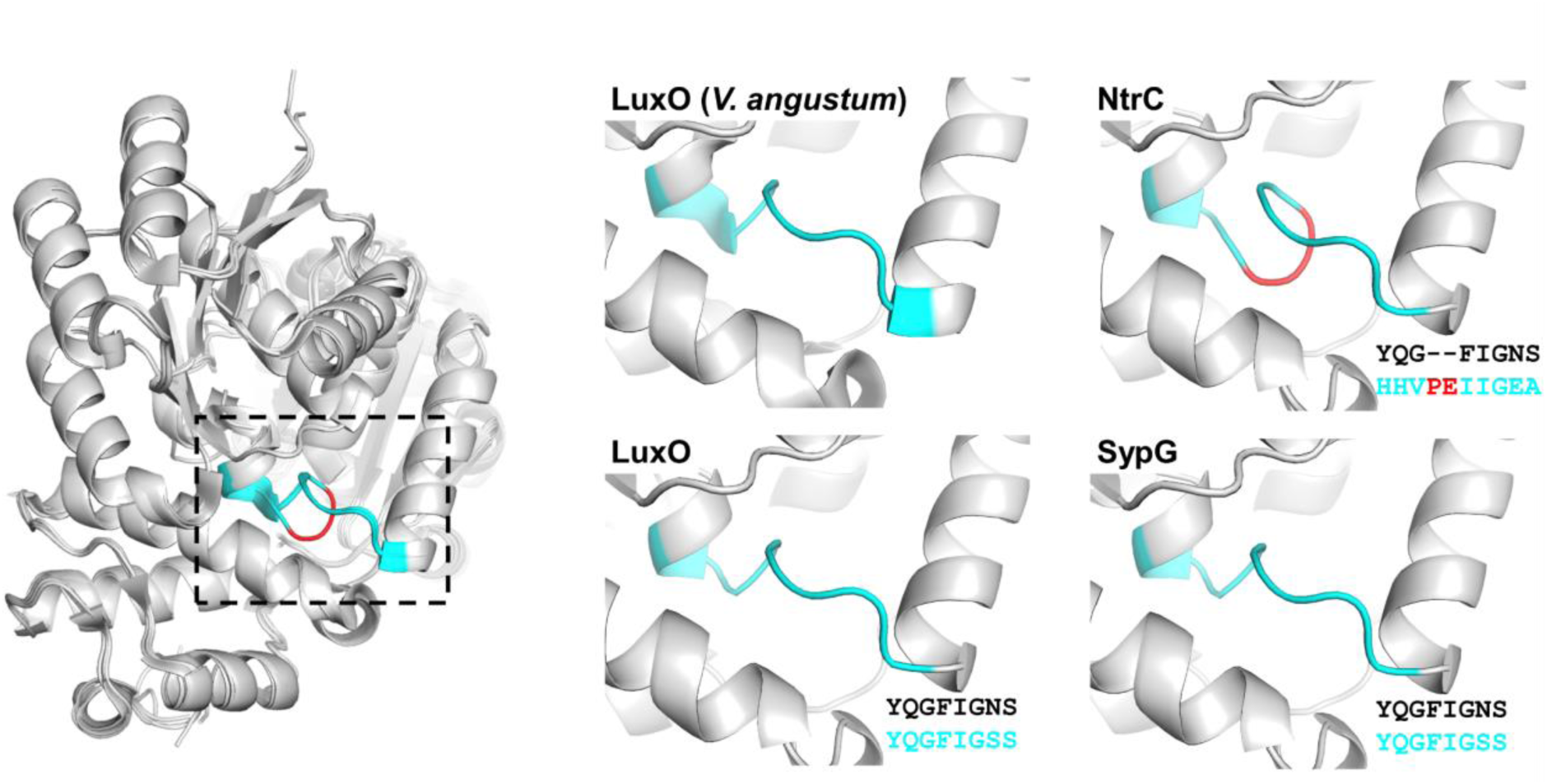
Alignment of SypG with LuxO crystal structure. *Left*, Overlay of homology models generated by MODELLER for the REC and AAA+ domains of LuxO (WP_011261589.1), SypG (WP_011263835.1), and NtrC (WP_011260965.1) with the crystal structure of the corresponding regions in the LuxO homolog of *V. angustum* (PDB 5EP0). Dotted box indicates the 8 residues of the linker region between REC and AAA+ domains (cyan). The deviation within the corresponding region of NtrC is shown in red. *Right*, the region indicated by the box for the individual models of the indicated bEBP. Sequences are the pairwise alignments of the 8 amino acids associated with the *V. angustum* reference (top) and indicated bEBP (bottom).

**Figure S3:**
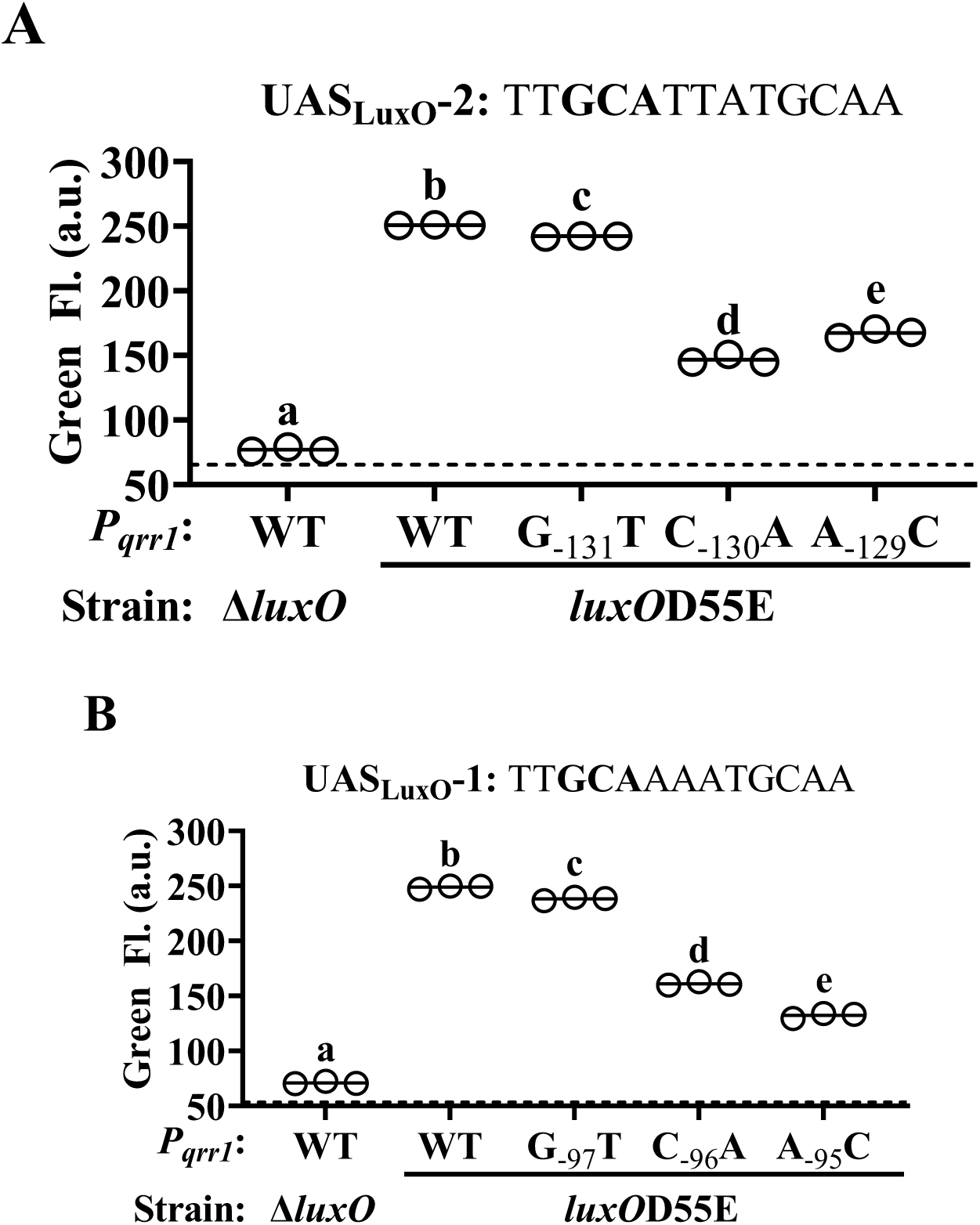
Transcriptional activation of *P_qrr1_* by LuxO. **(A)** Green fluorescence levels of TIM306 (Δ*luxO*) and CL59 (*luxO*D55E) harboring *P_qrr1_*::*gfp* reporter plasmids pEDR003, pEDS007 (G_-97_T), pEDS008 (C_-96_A), and pEDS009 (A_-95_C). The *luxO*D55E encodes a variant of LuxO that exhibits high activity in colonies. Sequence corresponds to 13-bp UAS_LuxO_-1, with nucleotides (-97)-(-95) that were individually mutated by site-directed mutagenesis shown in bold. Dotted line = autofluorescence cutoff. One-way ANOVA (F_4,10_ = 2998, *p* < 0.0001). **(B)** Green fluorescence levels of TIM306 (Δ*luxO*) and CL59 (*luxO*D55E) harboring *P_qrr1_*::*gfp* reporter plasmids pEDR003, pEDS007 (G_-97_T), pEDS008 (C_-96_A), and pEDS009 (A_-95_C). The *luxO*D55E encodes a variant of LuxO that exhibits high activity in colonies. Sequence corresponds to 13-bp UAS_LuxO_-1, with nucleotides (-97)-(-95) that were individually mutated by site-directed mutagenesis shown in bold. Dotted line = autofluorescence cutoff. One-way ANOVA (F_4,10_ = 5665, *p* < 0.0001).

**Figure S4:**
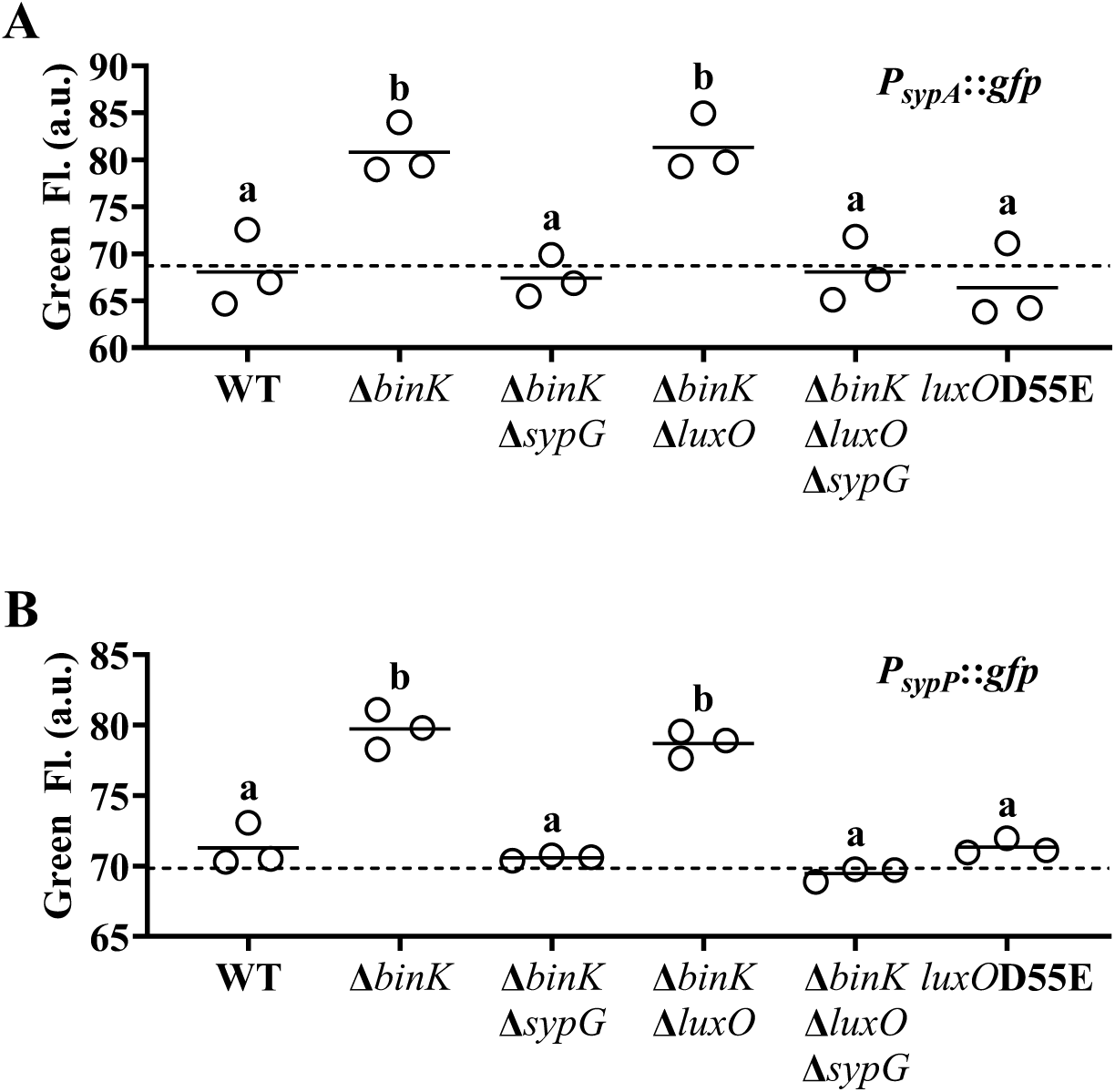
LuxO does not activate promoters of the *syp* locus in *V. fischeri*. **(A)** Green fluorescence levels of ES114 (WT), MJM2251 (Δ*binK*), EDR014 (Δ*binK* Δ*sypG*), EDR009 (Δ*binK* Δ*luxO*), EDR013 (Δ*binK* Δ*luxO* Δ*sypG*), and CL59 (*luxO*D55E) harboring *P_sypA_*::*gfp* reporter plasmid pVF_A1020P. Dotted line = autofluorescence cutoff. One-way ANOVA (F_5,12_ = 13.20, *p* = 0.0002). **(B)** Green fluorescence levels of ES114 (WT), MJM2251 (Δ*binK*), EDR014 (Δ*binK* Δ*sypG*), EDR009 (Δ*binK* Δ*luxO*), EDR013 (Δ*binK* Δ*luxO* Δ*sypG*), and CL59 (*luxO*D55E) harboring *P_sypP_*::*gfp* reporter plasmid pVF_A1035P. Dotted line = autofluorescence cutoff. One-way ANOVA (F_5,12_ = 61.10, *p* < 0.0001).

**Fig. S5:**
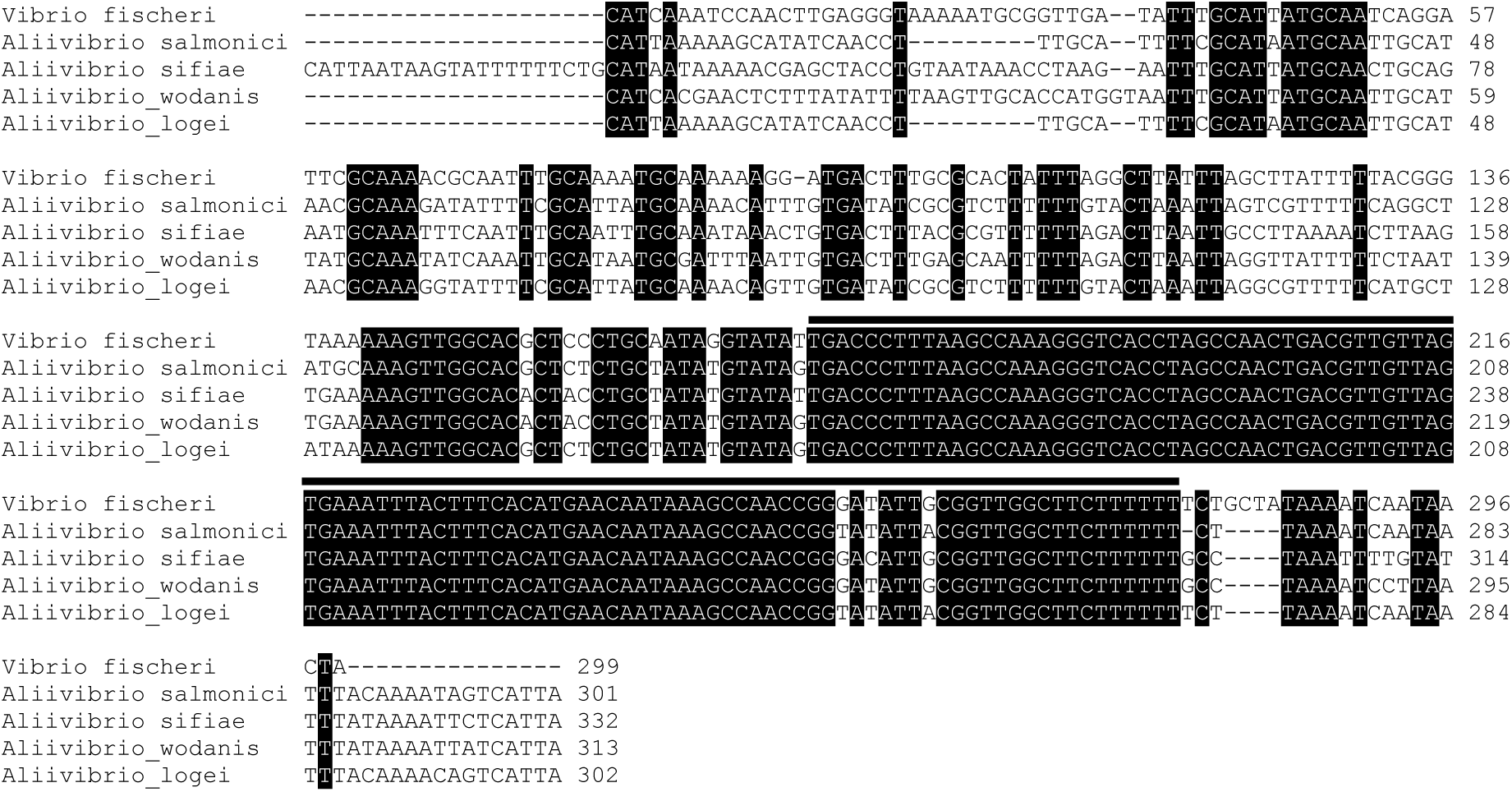
Conservation of Qrr1 within the Fischeri clade. Region between *uvrB* and *luxO* genes in indicated taxa were aligned by MEGA X. Highlighted positions indicate 100% identity across sequences. Line above alignment indicates positions predicted to encode a Qrr1 homolog. Accessions and regions analyzed: *Vibrio fischeri* = NC_006840.2 [complement(1033214..1033512)], *Aliivibrio salmonicida* = NC_011312.1 [complement(2009689..2009989)], *Aliivibrio sifiae* = NZ_MSCP01000001.1 [1631724..1632055], *Aliivibrio wodanis* = LN554846.1 [complement(1083567..1083879)], and *Aliivibrio logei* = NZ_MAJU01000008.1 [810639..810940].

**Figure S6:**
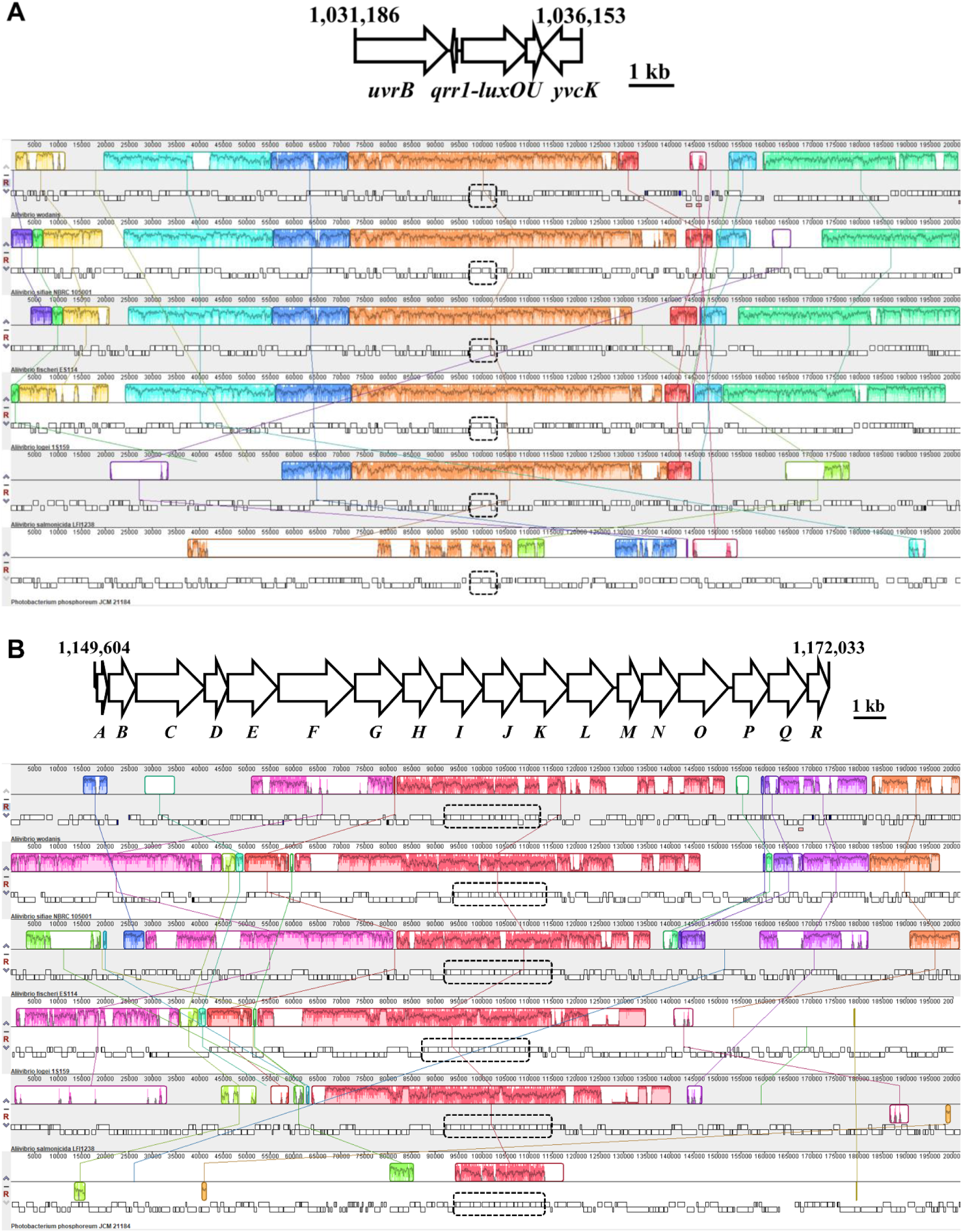
Conservation of gene synteny for *luxO* and *sypG* within the Fisheri clade. **(A)** Gene synteny associated with *luxO* in the Fisheri clade. *Top*, Region featuring *luxO* on chromosome 1 (NC_006840.2) of *V. fischeri* strain ES114. *Bottom*, ProgressiveMauve analysis of gene synteny associated with *luxO* for Fischeri clade members. Outlined regions indicate genomic blocks that are present in other taxa, with the interior plot indicating nucleotide similarity. *Photobacterium phosphoreum* was included as an outgroup. Accessions and regions analyzed: LN554846.1 [983877..1185307], NZ_MSCP01000001.1 [complement(1530329..1731726)], NC_006840.2 [933510..1134940], NZ_MAJU01000008.1 [complement(709196..910641)], NC_011312.1 [1909987..2111432], and NZ_MSCQ01000001.1 [complement(1671280..1872728)]. **(B)** Gene synteny associated with *sypG* in the Fisheri clade. *Top*, Region featuring *sypG* on chromosome 2 (NC_006841.2) of *V. fischeri* strain ES114. *Bottom*, ProgressiveMauve analysis of gene synteny associated with *sypG* for Fischeri clade members. Outlined regions indicate genomic blocks that are present in other taxa, with the interior plot indicating nucleotide similarity. *Photobacterium phosphoreum* was included as an outgroup. Accessions and regions analyzed: LN554847.1 [1266858..1468330], NZ_MSCP01000002.1 [1390955..1592439], NC_006841.2 [1057508..1259010], NZ_MAJU01000009.1 [(1..200000], NC_011313.1 [complement(241212..442708)], and NZ_MSCQ01000001.1 [complement(845650..1047125)].

**Figure S7:**
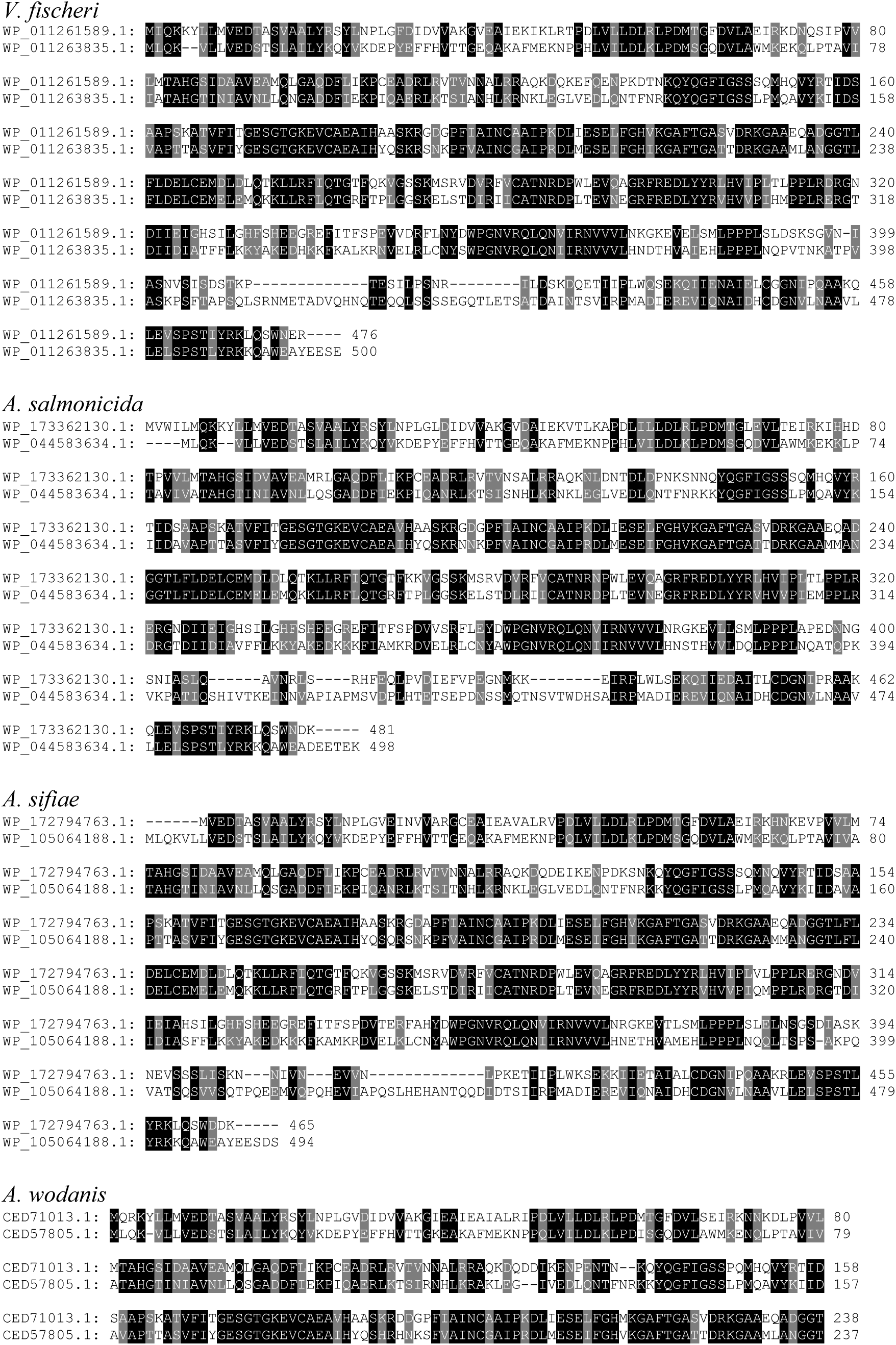
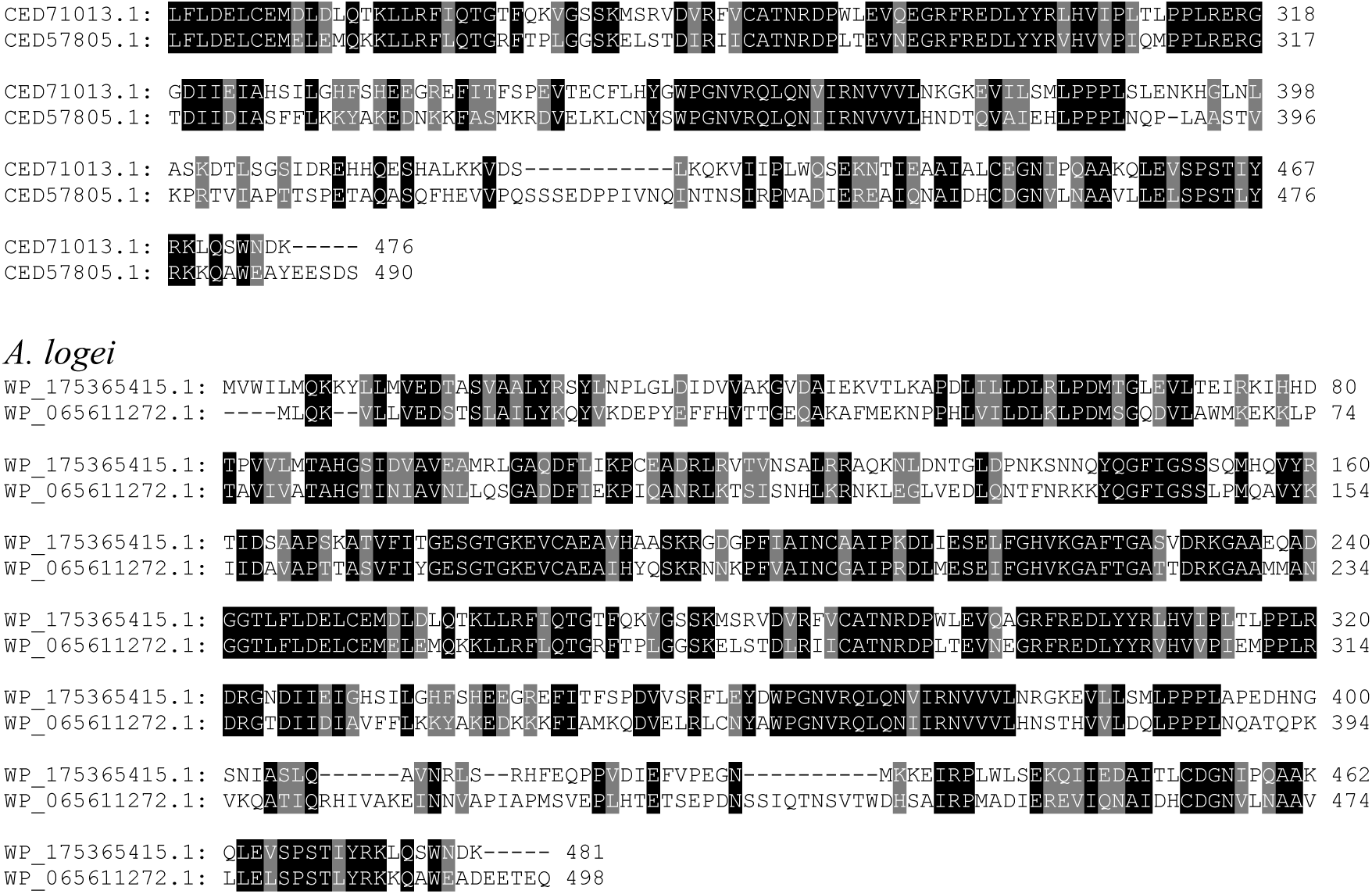
Pairwise alignments of LuxO (top sequence) and SypG (bottom sequence) homologs encoded by indicated Fischeri clade members. Amino acids highlighted in black (gray) indicate identity (similarity).

**Fig. S8:**
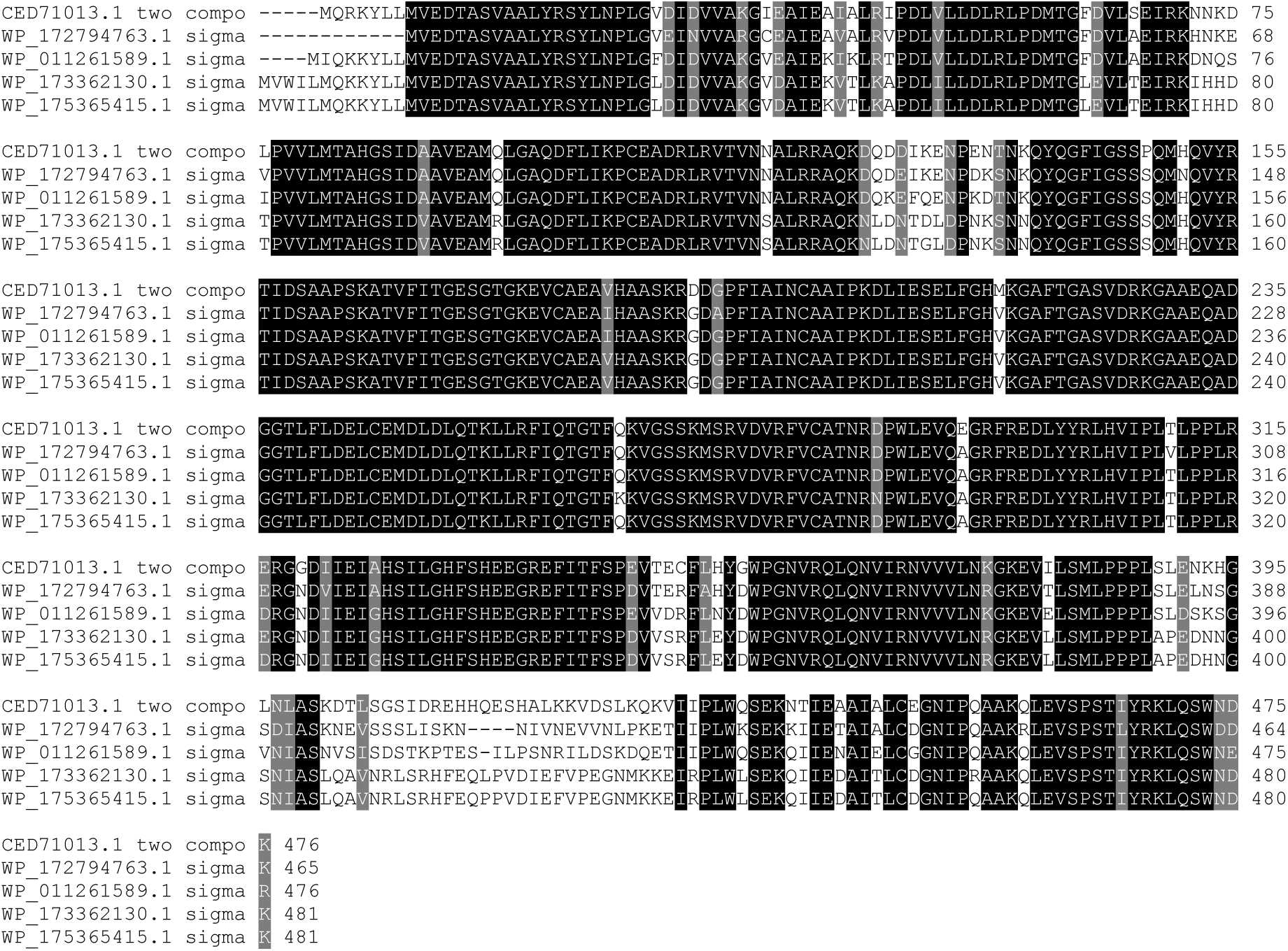
Multiple sequence alignment of LuxO homologs encoded by Fischeri clade members. Amino acids highlighted in black or gray indicate identity and similarity, respectively, among 100% of sequences. Top to bottom: *A. wodanis*, *A. sifiae*, *V. fischeri*, *A. logei*, and *A. salmonicida*.

**Figure S9:**
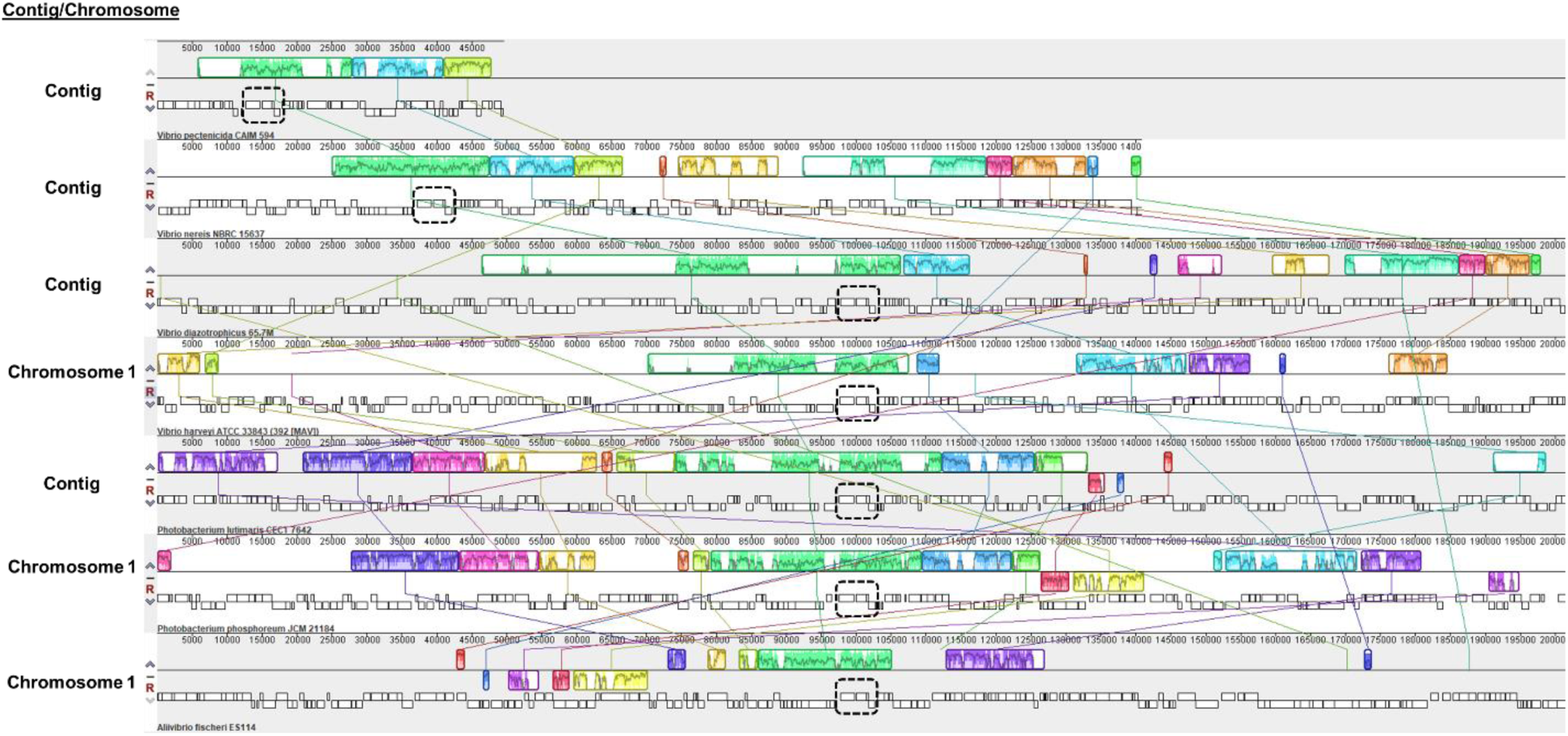
Gene synteny associated with *luxO* in *Vibrionaceae* taxa with incomplete genomes. Analysis of gene synteny by progressiveMauve for *luxO* encoded by indicated *Vibrionaceae* members. Outlined regions indicate genomic blocks that are present in other taxa, with the interior plot indicating nucleotide similarity. The gene cluster spanning *uvrB-yvcK* that contains *luxO* is indicated by the dashed outline. The regions linked to *luxO* for the fully assembled genomes of *V. harveyi*, *P. phosphoreum*, and *V. fischeri* are included for comparisons. Accessions and regions analyzed: NZ_RSFA01000020.1 [1..49638], NZ_BCUD01000001.1 [1..140875], NZ_POSL01000002.1 [complement(144684..346084)], NZ_CP009467.1 [complement(2563561..2764964)], NZ_SNZO01000002.1 [complement(542785..744227)], NZ_MSCQ01000001.1 [complement(1671280..1872728)], and NZ_MSCQ01000001.1 [complement(1671280..1872728)].

**Fig. S10:**
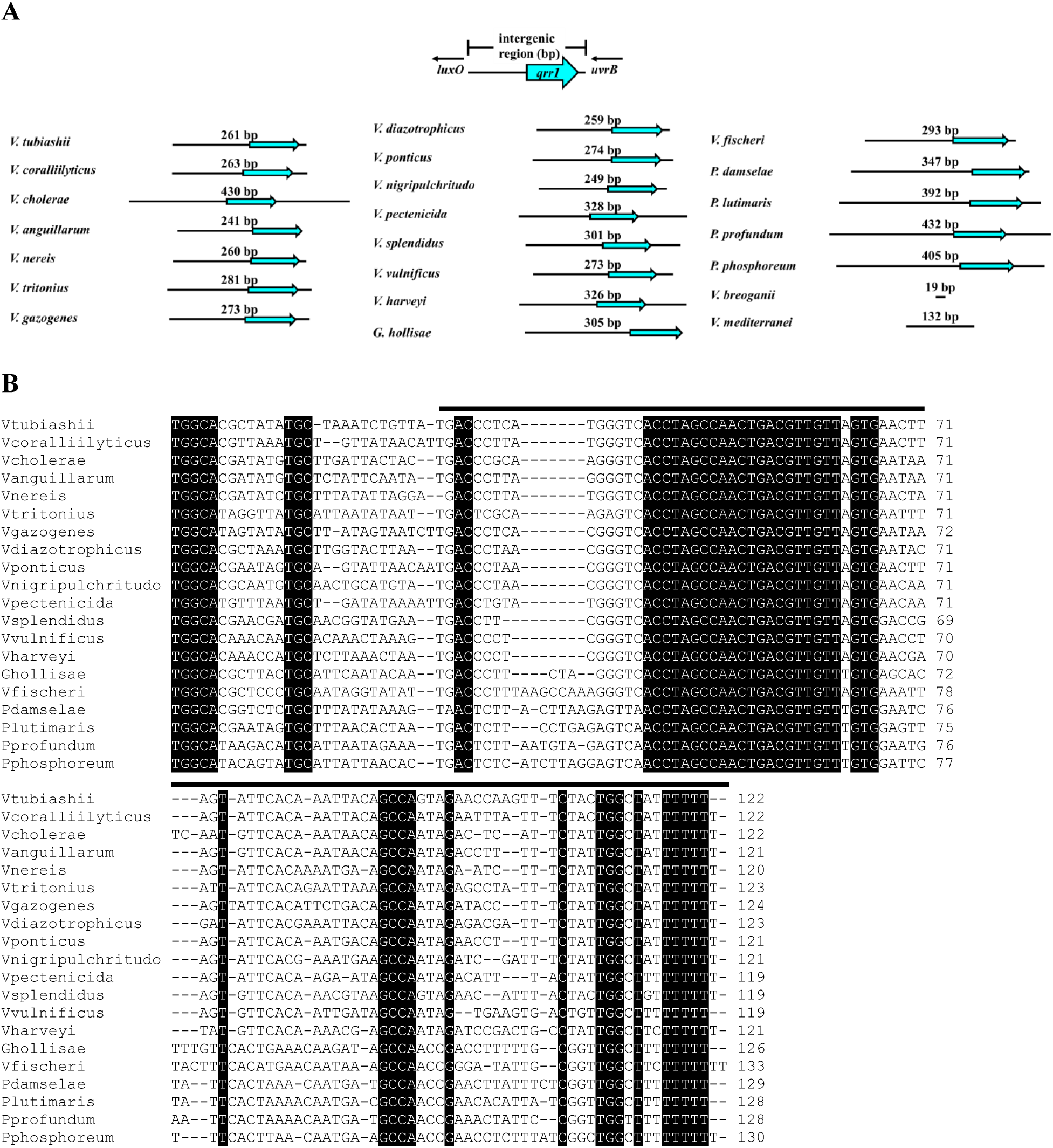
Conservation of *qrr* gene within *uvrB*-*luxO* intergenic region among *Vibrionaceae*. **(A)** Cartoon indicating length in bp of *uvrB*-*luxO* intergenic region for indicated taxa. Location of putative gene encoding a Qrr is denoted by the cyan arrow. **(B)** Sequences of homologs of Qrr1 (denoted by the black lines) encoded within the *uvrB*-*luxO* intergenic region of each indicated taxon. Each homolog was identified by first using the locations of the -24 (GGC) and -12 (GC) sites corresponding to σ^54^ binding sites to determine the putative transcriptional start site and then locating the thymidine repeat corresponding to the likely terminator sequence.

**Fig. S11:**
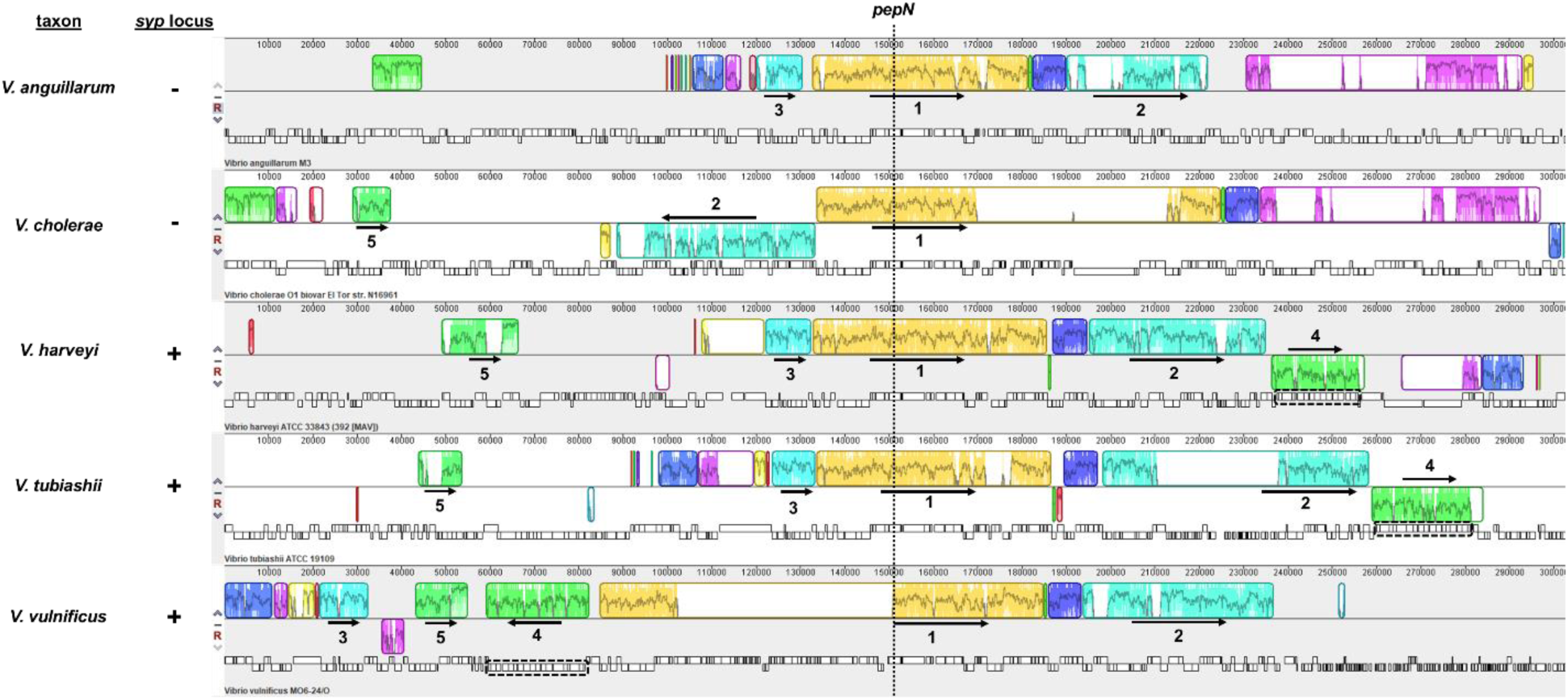
Gene synteny associated with *pepN* among select *Vibrionaceae*. Analysis of gene synteny by progressiveMauve for PepN homolog (identified by BLAST using WP_069687131.1) encoded by indicated *Vibrionaceae* members. Dotted line indicates *pepN* gene. Outlined regions indicate locally colinear blocks (LCBs) that are present in at least two taxa, with the interior plot indicating nucleotide similarity. Numbers identify the LCBs that were used to interpret gene synteny, with the arrows indicating the relative LCB orientation. The *syp* locus is indicated by the dashed outline and associated with LCB 4. *V. harveyi* and *V. tubiashii* feature the *syp* locus downstream of LCB 2. In contrast, the *syp* locus of *V. vulnificus* is in an opposite orientation and located between LCBs 3 and 1. Neither *V. anguillarum* nor *V. cholerae* feature a *syp* locus. Accessions and regions analyzed: NC_022223.1 [complement(1659437..1962043)], NC_002505.1 [complement(1452881..1755487)], NZ_CP009467.1 [complement(1843797..2146403)], NZ_CP009354.1 [1461021..1763627], and NC_014965.1 [complement(1499333..1801960)].

**Fig. S12:**
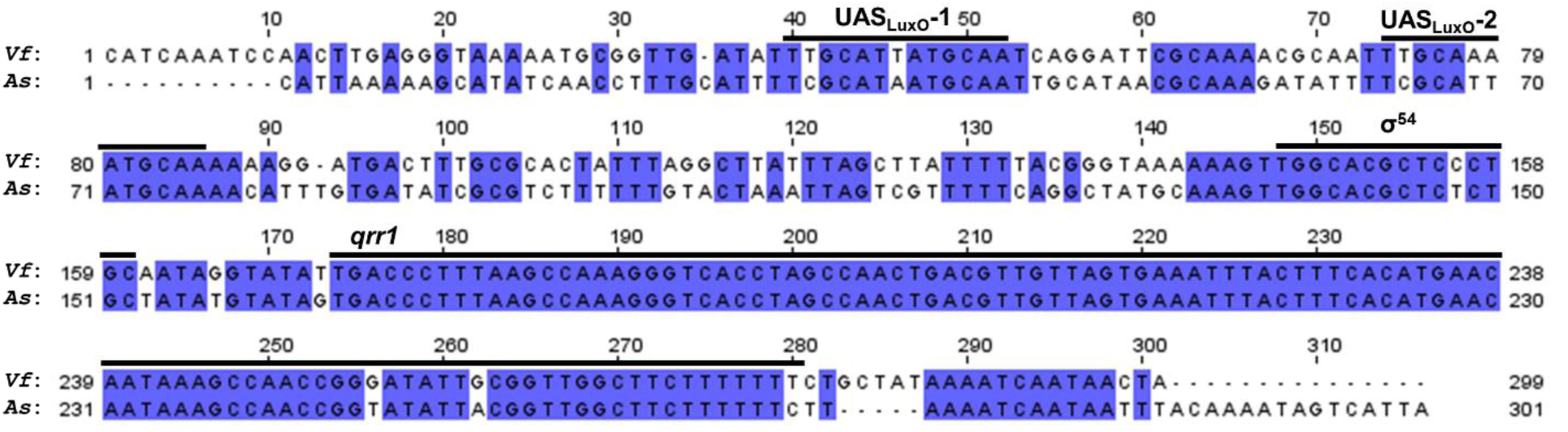
Conservation of the *qrr1* locus among *V. fischeri* and *A. salmonicida*. Pairwise alignment of *luxO*-*uvrB* intergenic region in *V. fischeri* (*Vf*) and *A. salmonicida* (*As*) generated by ClustalW in MEGA X, with identical nucleotides highlighted. Regions relevant to the expression of Qrr1 are labeled.

